# Mechanosensitive activation of mTORC1 mediates ventilator induced lung injury during the acute respiratory distress syndrome

**DOI:** 10.1101/2020.03.02.973081

**Authors:** Hyunwook Lee, Qinqin Fei, Adam Streicher, Wenjuan Zhang, Colleen Isabelle, Pragi Patel, Hilaire C. Lam, Miguel Pinilla-Vera, Diana Amador-Munoz, Diana Barragan-Bradford, Angelica Higuera, Rachel K. Putman, Elizabeth P. Henske, Christopher M. Bobba, Natalia Higuita-Castro, R. Duncan Hite, John W. Christman, Samir N. Ghadiali, Rebecca M. Baron, Joshua A. Englert

**Author notes:** denotes equal contribution (Rebecca M. Baron, Joshua A. Englert). **Corresponding author:** Joshua A. Englert, MD, Division of Pulmonary, Critical Care, & Sleep Medicine, The Ohio State University Wexner Medical Center, 201 Davis Heart and Lung Research Institute, 473 West 12th Avenue, Columbus, OH 43210.

## Abstract

Acute respiratory distress syndrome (ARDS) is a highly lethal condition that impairs lung function and causes respiratory failure. Mechanical ventilation maintains gas exchange in patients with ARDS, but exposes lung cells to physical forces that exacerbate lung injury. Our data demonstrate that mTOR complex 1 (mTORC1) is a mechanosensor in lung epithelial cells and that activation of this pathway during mechanical ventilation exacerbates lung injury. We found that mTORC1 is activated in lung epithelial cells following volutrauma and atelectrauma in mice and humanized in vitro models of the lung microenvironment. mTORC1 is also activated in lung tissue of mechanically ventilated patients with ARDS. Deletion of *Tsc2*, a negative regulator of mTORC1, in epithelial cells exacerbates physiologic lung dysfunction during mechanical ventilation. Conversely, treatment with rapamycin at the time mechanical ventilation is initiated prevents physiologic lung injury (i.e. decreased compliance) without altering lung inflammation or barrier permeability. mTORC1 inhibition mitigates physiologic lung injury by preventing surfactant dysfunction during mechanical ventilation. Our data demonstrate that in contrast to canonical mTORC1 activation under favorable growth conditions, activation of mTORC1 during mechanical ventilation exacerbates lung injury and inhibition of this pathway may be a novel therapeutic target to mitigate ventilator induced lung injury during ARDS.

## Introduction

The acute respiratory distress syndrome (ARDS) is a devastating condition that affects over 200,000 patients annually in the U.S. with mortality rates up to 45% in its most severe form.(1–3) The only treatment for patients with ARDS is supportive care with mechanical ventilation (MV), and although life-saving, MV can exacerbate pre-existing lung injury and even cause *de novo* injury, known as ventilator induced lung injury (VILI).(4) Limiting lung distention by using low tidal volume (TV) ventilation decreases mortality in ARDS patients,(5) but factors such as regional heterogeneity lead to persistent injury even with low tidal volume ventilation.(6) VILI arises from three injurious forces including excessive stretch (volutrauma), increased transmural pressure (barotrauma), and mechanical stress from repetitive collapse and re-opening of lung units (atelectrauma).(7) The molecular mechanisms by which these injurious forces cause lung injury remain poorly understood which has limited the development of targeted therapies to prevent lung injury in patients requiring mechanical ventilation and in patients with ARDS.

Mechanotransduction is the process by which physical forces are transduced into biologic responses. The lungs stretch cyclically during spontaneous breathing and positive pressure mechanical ventilation. The precise mechanisms by which the lungs sense and respond to injurious forces during mechanical ventilation are not well understood. Several cell types have been implicated in the transduction of mechanical signals following injurious ventilation including epithelial(8, 9) and endothelial cells (10). One approach to elucidate the molecular mechanisms of lung injury during mechanical ventilation is to identify mechanosensitive signaling pathways that are differentially activated by the various types of VILI.

mTOR complex 1 (mTORC1) is a ubiquitously expressed multi-protein complex with serine/threonine kinase activity that is involved in cellular metabolism and response to stress. mTORC1 activation has been also been implicated in mechanotransduction in muscle.(11–13) In the lung, mTORC1 contributes to the pathogenesis of lung injury in response to cigarette smoke (14) and endotoxin (15). However the role of mTORC1 activation in response to mechanical forces in the lung and its role in the pathogenesis of VILI has not been examined. In contrast to canonical activation of mTORC1 under favorable growth conditions, we discovered that mTORC1 is pathologically activated by injurious forces during mechanical ventilation. Based on this observation, we investigated whether mTORC1 activation mediates the development of lung injury following injurious mechanical ventilation and whether inhibition of this pathway might represent a novel therapeutic target for patients with ARDS.

## Results

### Volutrauma and atelectrauma activate mTORC1 in epithelial cells in murine models of injurious mechanical ventilation

We used a model of simultaneous volutrauma (tidal volume (V_T_, 12 cc/kg) and atelectrauma (positive end expiratory pressure [PEEP] 0 cm H_2_O) that impaired lung function, induced inflammation (i.e. bronchoalveolar lavage (BAL) neutrophils and IL6 levels), and increased barrier permeability (i.e. BAL protein levels) **(Supplemental Figure 1).** mTORC1 activation was assessed in lung tissue by immunoblotting for phosphorylated isoforms of the ribosomal S6 protein (16, 17) and S6 kinase.(18) Spontaneously breathing control mice and mice ventilated with non-injurious settings (V_T_ 6 cc/kg, PEEP 5 cm H_2_O) had low levels of mTORC1 activation. In contrast, mice ventilated with injurious high tidal volumes (12 cc/kg) or without PEEP had increased levels of phosphorylated S6 **(Figure 1A)**. Immunostaining revealed that the primary site of mTORC1 activation was the epithelium, with the most intense activation in the airway epithelium **(Figure 1B-D)**. Given that critically ill patients frequently develop ARDS and require mechanical ventilation (MV) in the setting of existing lung injury, we also subjected mice to a 2-hit model of acute respiratory distress syndrome (ARDS) in which MV was initiated in the setting of polymicrobial sepsis induced by cecal ligation and puncture (CLP). Twenty-four hours after sub-lethal CLP or sham operation, mice were subjected to injurious MV (CLP/VILI-12 cc/kg, PEEP=2.5) for 4 hours. Mice subjected to CLP/VILI had impaired lung function and increased markers of lung injury compared to uninjured control mice or either injury alone **(Supplemental Figure 2)** which correlated with increased mTORC1 activation in whole lung tissue **(Figure 1E)**. As was the case with VILI alone, the most prominent site of mTORC1 activation following CLP/VILI was the epithelium. These data demonstrate that mTORC1 is activated in lung epithelial cells following injurious mechanical ventilation.

**Figure 1.**
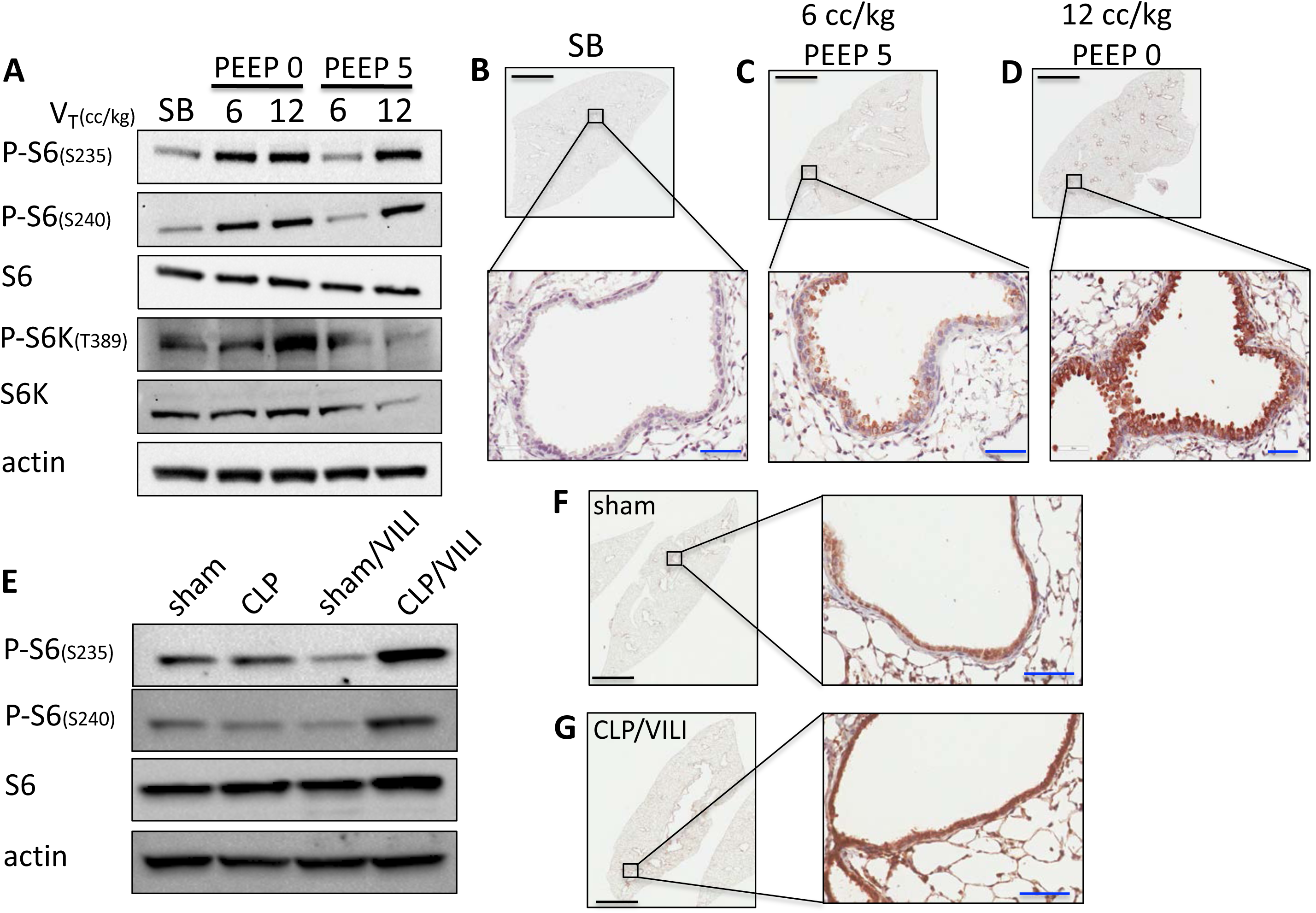
Injurious mechanical ventilation activates mTORC1 in lung epithelial cells. A) Immunoblots of phosphorylated and total ribosomal S6, S6 kinase (S6K), and beta-actin using pooled protein lysate from whole lung tissue of spontaneously breathing (SB) control mice (n=3/lane) or mice subjected to mechanical ventilation (n=4/lane) with high tidal volume (V_T_, 12 cc/kg), low tidal volume (V_T_ 6 cc/kg) with or without the use of positive end expiratory pressure (PEEP, cm H_2_O). Low power (4X) and high power (400X, inset) images from lung tissue that was immunostained for phosphorylated S6 (P-S6, S235) from SB control mice (B), mice ventilated with non-injurious settings (C, V_T_ 6 cc/kg, PEEP 5 cm H_2_O), and mice ventilated with injurious settings (D, V_T_ 12 cc/kg, PEEP 0 cm H_2_O). E) Immunoblots from whole lung tissue of mice subjected to sham laparotomy (n=3/lane), cecal ligation and puncture (CLP, n=4/lane), ventilator induced lung injury (VILI-V_T_ 12 cc/kg, PEEP 2.5 cm H_2_O) 24 hrs after sham laparotomy (n=3/lane), and VILI 24 hrs after CLP (CLP/VILI, n=4/lane). Representative images from lung tissue that was immunostained for P-S6 (S235) from sham (F) and CLP/VILI (G) mice. black bars=2 mm, blue bars= 50 μm

### Tsc2 deletion in airway epithelial cells exacerbates lung injury during mechanical ventilation

To determine whether mTORC1 activation in lung epithelial cells mediates the development of lung injury or represents a compensatory response that promotes recovery, we used a genetic approach to generate mice with increased mTORC1 in airway epithelial cells. We bred mice with homozygously floxed alleles of *Tsc2* (a negative regulator of mTORC1) (19) to mice expressing Cre recombinase under the control of the airway epithelial specific CC10 promoter (20). CC10-Cre mice were bred to mT/mG reporter mice (21) to confirm airway epithelial specific expression of the Cre recombinase prior to breeding with *Tsc2* flox mice. (**Supplemental Figure 3**).

*CC10*^Cre^/*Tsc2*^flox/flox^ progeny were backcrossed to a C57BL/6 background and examination of lung morphology and function revealed no abnormalities at baseline (**Supplemental Figure 3**). There were no differences in the number of BAL cells (**Supplemental Figure 3**) and differential counting revealed that >95% of cells were alveolar macrophages (data not shown). Low levels of mTORC1 activation (i.e. P-S6) were observed in spontaneously breathing mice Cre-control. Mice with airway epithelial *Tsc2* deletion (Cre+) mice had increased mTORC1 activation in airway epithelial cells by immunohistochemical staining. **(Supplemental Figure 3)** Primary tracheobronchial epithelial cells (22) isolated from Cre+ mice had decreased tuberin (encoded by *Tsc2)* expression. **(Supplemental Figure 3).** Cre+ mice were subjected to simultaneous volutrauma and atelectrauma (12 cc/kg, PEEP 0 cm H_2_O) for 4 hours and had increased lung stiffness (i.e. elastance) and decreased inspiratory capacity compared to Cre-controls **(Figure 2A-B)**. Cre+ mice were also more hypoxemic during injurious ventilation **(Figure 2C)**. Notably, recruitment of inflammatory cells, barrier permeability (BAL protein), and pro-inflammatory BAL cytokine and chemokine levels were not different following VILI or in spontaneously breathing control mice **(Figure 2D-H, Supplemental Figure 4)**. These data demonstrate that airway epithelial *Tsc2* deletion exacerbates physiologic lung injury (i.e. increases lung stiffness, decreases inspiratory capacity, and impairs oxygenation) following injurious ventilation without altering lung inflammation or barrier dysfunction.

**Figure 2.**
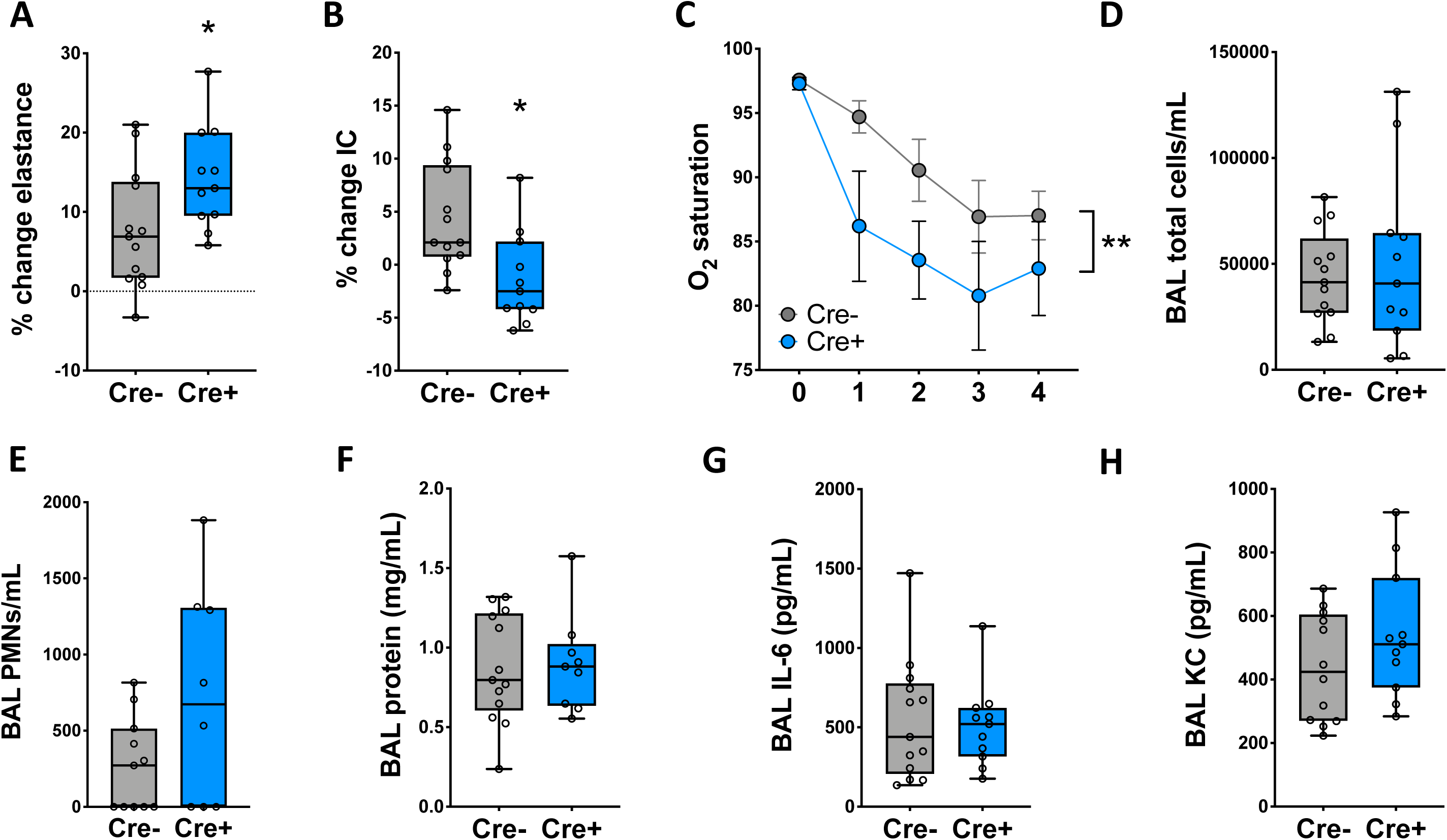
Airway epithelial *Tsc2* deletion exacerbates lung injury in a murine model of combined volutrauma and atelectrauma. Mice with airway epithelial *Tsc2* deletion (Cre+) and Cre-control mice were subjected to mechanical ventilation with high tidal volume (12 cc/kg) without positive end expiratory pressure (PEEP 0 cm H_2_O) for 4 hrs. The change in lung elastance (A) and inspiratory capacity (B, IC) were measured during mechanical ventilation. All data were normally distributed, analyzed by student’s t-test, *p<0.05 C) Oxygen saturations were measured via pulse oximetry during the 4 hr period of ventilation. **indicates that genotype is a statistically significant factor, p<0.05, by 2-way ANOVA with repeated measures with Holms-Sidak post hoc test. Following mechanical, a bronchoalveolar lavage (BAL) was performed and total inflammatory cells (D), neutrophils (E, PMNs), and protein levels (F) were measured. For PMN counts, n=12 for Cre- and n=8 for Cre+. IL-6 (G) and KC (H) levels were measured in BAL fluid by ELISA. n=13 for Cre- and n=11 for Cre+ unless otherwise noted.

### Mechanically ventilated patients with diffuse alveolar damage have increased mTORC1 activation in lung tissue

To assess whether mTORC1 is activated in patients with ARDS undergoing mechanical ventilation, we obtained formalin-fixed lung tissue from a pathology tissue bank. Specimens were analyzed from 5 patients with diffuse alveolar damage (DAD), the most common lung pathology in ARDS patients.(23, 24) Normal lung tissue adjacent to resected lung tumors from 5 subjects was used as a control. Clinical characteristics of the subjects are shown in **Supplemental Table 1** and there were no significant differences between groups. Notably all patients were receiving mechanical ventilation at the time of biopsy. Immunostaining for phosphorylated ribosomal S6 revealed increased mTORC1 activation in lung tissue from patients with DAD compared to normal lung tissue. **(Figure 3A-F, Supplemental Figure 5)** Although diffuse staining was present throughout the lung, it appeared that the most prominent site of staining was the epithelium including airway epithelial cells. **(Figure 3D-F)** The intensity of P-S6 staining was quantitated by image analysis and was significantly higher in patients with DAD compared to controls. **(Figure 3G)** To determine whether mTORC1 activation was due to the underlying lung injury or the use of mechanical ventilation, we obtained additional biopsies from spontaneously breathing patients undergoing transbronchial biopsies during bronchoscopy or from surgical wedge resections from mechanically ventilated patients. Biopsies had a variety of pathologic findings including several with diffuse alveolar damage, several types of interstitial lung disease, airway diseases, and no specific pathology. In general, spontaneously breathing patients had less mTORC1 activation compared to mechanically ventilated patients regardless of the underlying pathology. **(Supplemental Figure 6)** Patients without lung injury had low levels of mTORC1 activation. **(Supplemental Figure 6)** These data suggest that injurious mechanical ventilation in the context of underlying lung injury activates mTORC1 in the lung epithelium.

**Figure 3.**
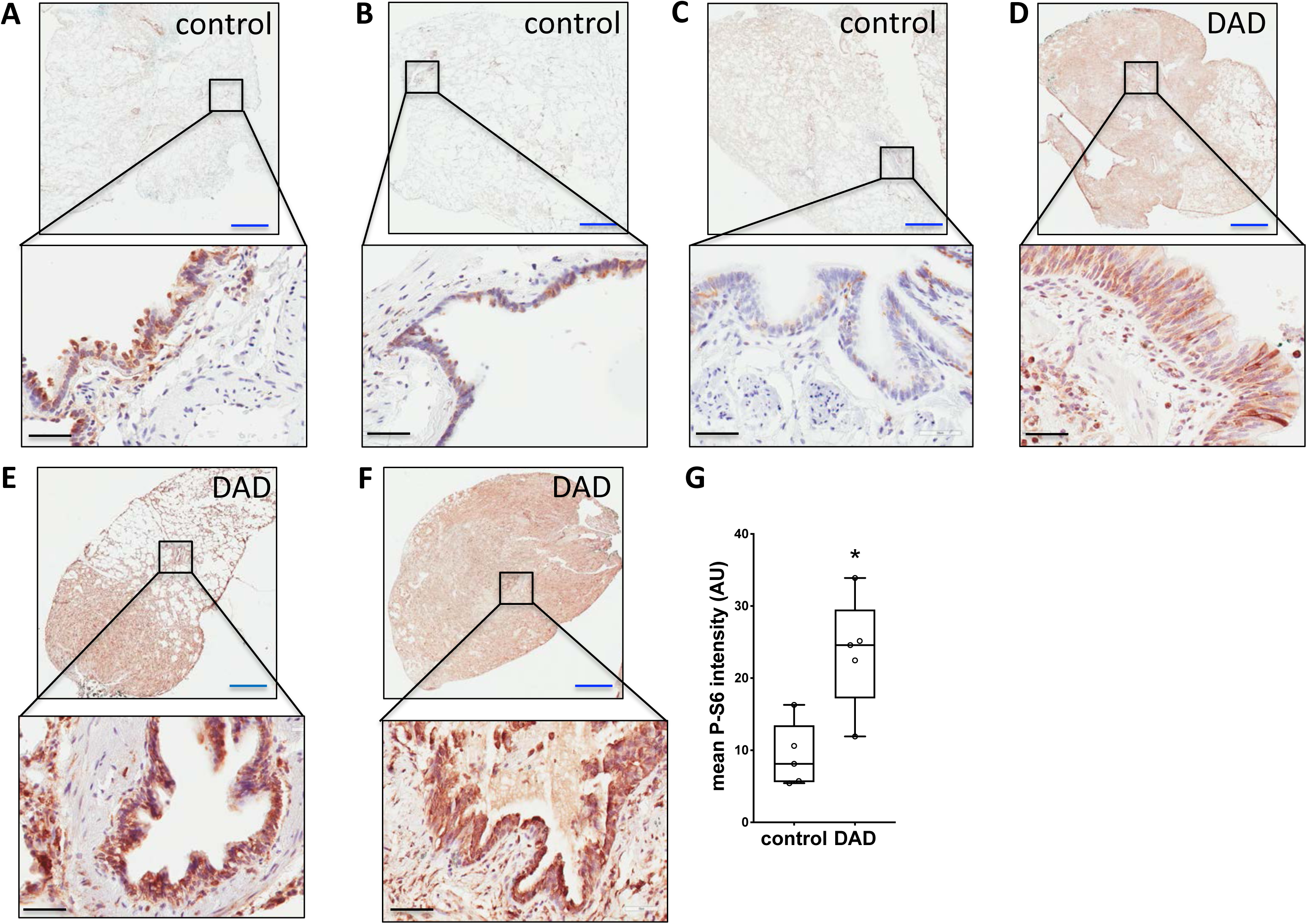
mTORC1 is activated in lung tissue from mechanically ventilated patients with diffuse alveolar damage. Photomicrographs of lung sections stained for P-S6 (Ser235/236) from normal lung tissue from control patients (A-C) and from patients with diffuse alveolar damage (DAD, D-F). Mean intensity of staining of the lung tissue was quantitated using low power images of the entire slide (n=5/group). Blue bar=2 mm and black bar=50 μm. *p<0.05 vs control by student’s t-test.

### In vitro models of volutrauma and atelectrauma activate mTORC1 in primary human airway epithelial cells

To investigate the molecular mechanisms of mTORC1 activation in epithelial cells following mechanical ventilation we utilized in vitro models of the various forms of ventilator induced lung injury **(Figure 4A-C)**. To simulate volutrauma, primary human airway epithelial cells were grown on flexible membranes and subjected to cyclic stretch that models overdistention during injurious mechanical ventilation (25, 26). Volutrauma increased phosphorylation of S6 kinase and ribosomal S6 within 30 minutes in distal small airway epithelial cells (SAECs) and proximal human bronchial epithelial cells (HBE) **(Figure 4D, Supplemental Figure 7)**. mTORC1 activation was dose and time dependent and rapidly decreased following cessation of injurious stretch. (**Supplemental Figure 7**). Injurious stretch also increased phosphorylation of 4E-BP1, a regulator of protein translation that is phosphorylated when mTORC1 is activated (27) **(Figure 4D)**. Volutrauma induced mTORC1 activation was prevented by allosteric (*i.e.* rapamycin) and active site (*i.e.* Torin 2) mTOR inhibitors **(Supplemental Figure 7)**. To model airway collapse and reopening during atelectrauma, we grew SAECs to confluence on collagen coated glass slides and exposed them to a moving air-liquid interface by propagating a fluid bubble over the monolayer which simulates the movement of edema fluid in ARDS patients undergoing mechanical ventilation.(28) As was seen with volutrauma, atelectrauma rapidly increased P-S6 levels in SAECs **(Figure 4E)**. Interestingly, we did not see the increase in S6 kinase or 4-EBP1 phosphorylation that was seen following volutrauma. To model barotrauma, SAECs were cultured at an air-liquid interface and cells were subjected to high levels of transmural pressure.(29, 30) Although mTORC1 was activated following barotrauma, the degree of activation was much less than that seen with volutrauma and atelectrauma. **(Figure 4F)**. In summary, these data demonstrate that mechanical forces during in vitro volutrauma and atelectrauma activate mTORC1 mirroring the findings from our murine model and patients with ARDS.

**Figure 4.**
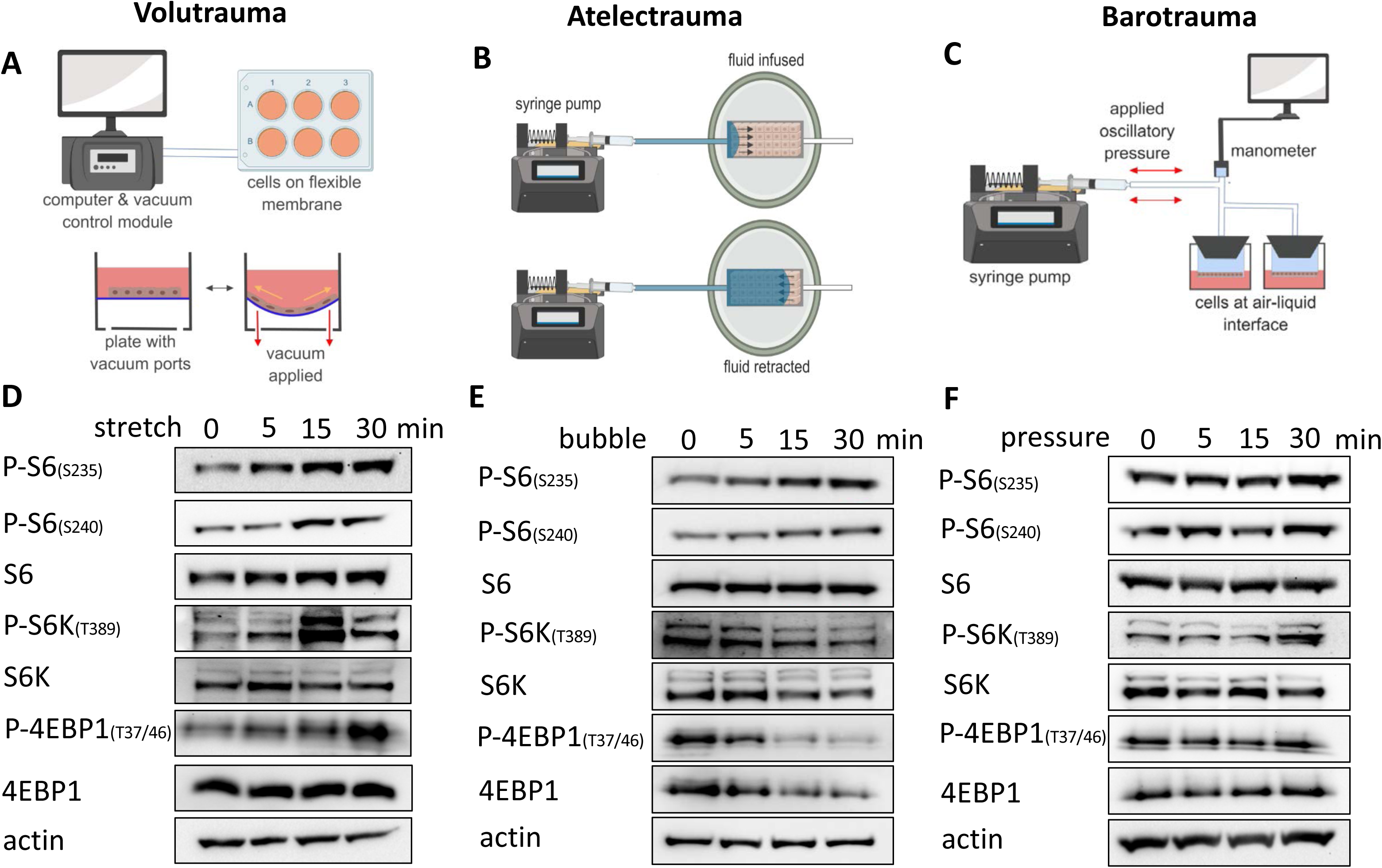
mTORC1 is rapidly activated by volutrauma and atelectrauma in vitro. Schematics of in vitro volutrauma, atelectrauma, and barotrauma models used. (A-C) D) Small airway epithelial cells (SAECs) were subjected to equibiaxial stretch (20%, 0.2 Hz) for varying amounts of time prior to immunoblotting for markers of mTORC1 activation (protein pooled from n=3 wells/lane). E) SAECs were grown to confluent monolayers on collagen coated glass slides in a microfluidic chamber and subjected to bubble flow (velocity 30 mm/sec) to model atelectrauma for varying amounts of time prior to immunoblotting for makers of mTORC1 activation (protein pooled from n=2 gels/timepoint) F) SAECs were grown at air-liquid interface on Transwells and subjected to oscillatory pressure (20 cm H_2_O, 0.2 Hz) for varying amounts of time prior to immunoblotting for markers of mTORC1 activation (protein pooled from n=3 wells/lane).

### Volutrauma activates mTORC1 through reactive oxygen species-dependent activation of the ERK pathway

We sought to determine the molecular mechanisms by which injurious ventilation activates mTORC1. Several canonical signaling pathways integrate biochemical and physiologic cues and activate mTORC1 via a series of phosphorylation events, including the ERK and Akt pathways. To determine whether these pathways mediate mTORC1 activation in our volutrauma model, we treated human primary airway epithelial cells with a specific ERK 1/2 (SCH772984) or Akt (MK-2206) inhibitor immediately prior to injurious stretch. In vitro volutrauma rapidly increased ERK1/2 phosphorylation that was potently inhibited by SCH772984 **(Figure 5A)**. ERK inhibition completely abrogated the increase in S6 phosphorylation following injurious stretch indicating that stretch induced mTORC1 activation is ERK dependent **(Figure 5A)**. In contrast to ERK activation by volutrauma, in vitro stretch did not increase Akt phosphorylation **(Figure 5B).** Treatment with MK-2206 potently inhibited basal Akt and mTORC1 activation, but did not prevent S6 phosphorylation during volutrauma indicating that VILI induced mTORC1 activation is Akt independent. **(Figure 5B)** A variety of second messengers can activate ERK1/2 in the lung epithelium including cellular reactive oxygen species. (31, 32) Specifically, hydrogen peroxide (H_2_O_2_) can activate ERK 1/2 and mTORC1 in certain cell types. (33–35) To determine if cellular ROS are involved in ERK-dependent mTORC1, airway epithelial cells were subjected to volutrauma in vitro and cellular ROS were measured. Total ROS increased following 30 minutes of volutrauma in airway epithelial cells **(Figure 5C-D)** which were, in part, from mitochondria **(Figure 5E-F)**. Treatment of airway epithelial cells with exogenous H_2_O_2_ increased phosphorylation of ERK1/2 within 5 minutes and preceded a dose dependent activation of mTORC1 that peaked between 30-60 minutes **(Figures 5G, Supplemental Figure 8)**. To determine if stretch induced mTORC1 activation was dependent on the release of cellular ROS, we treated SAECs with glutathione ethyl ester (GSH-EE), a cell permeable form of cellular antioxidant glutathione (36), and found that stretch induced mTORC1 activation was completed abrogated **(Figure 5H)**. These data indicate that the mechanism of mTORC1 activation during volutrauma involves the release of cellular ROS that act as second messengers to activate mTORC1 via the canonical ERK pathway.

**Figure 5.**
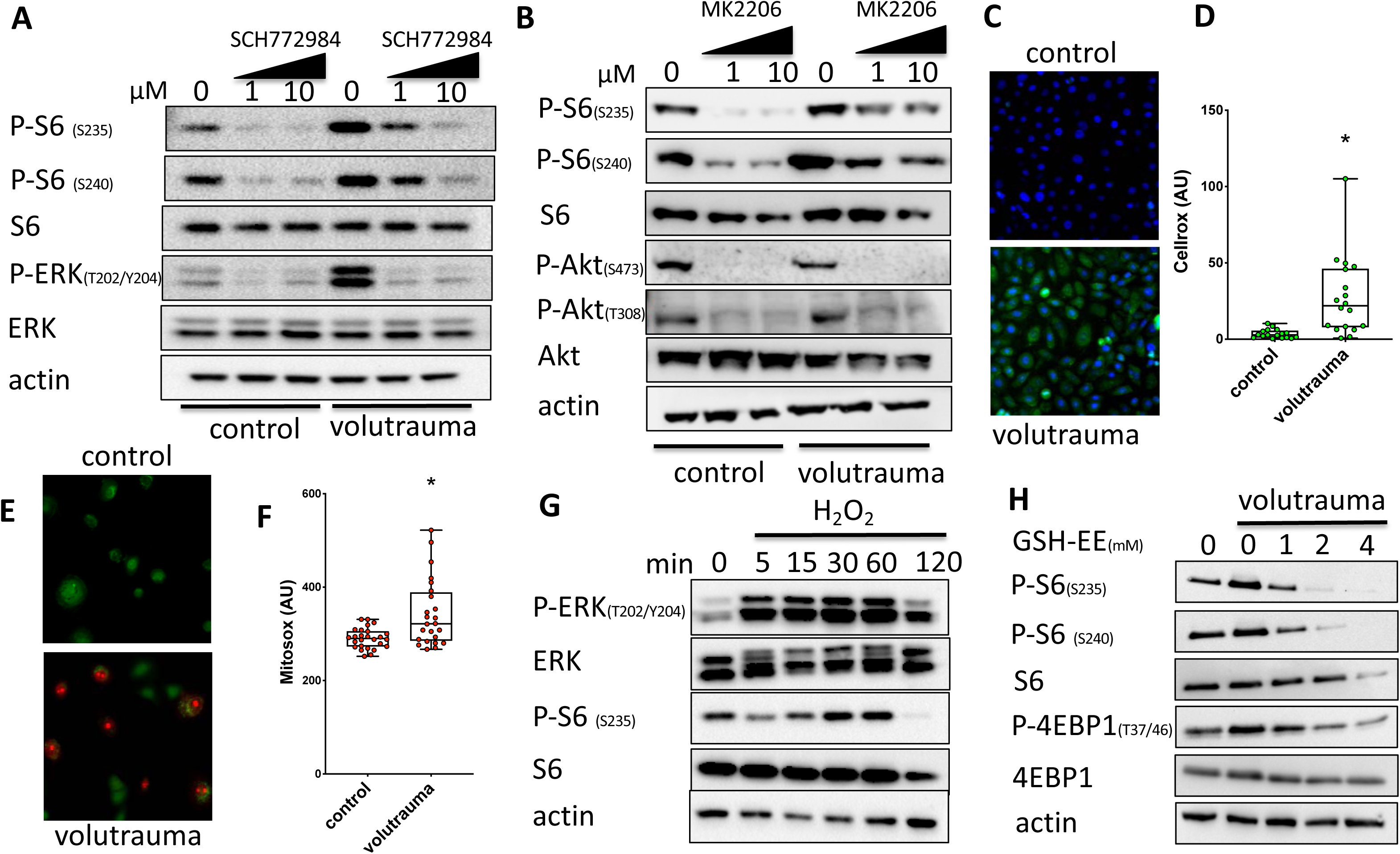
In vitro volutrauma activates mTORC1 through reactive oxygen species-dependent activation of the ERK pathway. A) Human bronchial epithelial cells were subjected to volutrauma (20% equibiaxial stretch, 0.2 Hz, 30 min) in the presence of increasing doses of the ERK 1/2 inhibitor (SCH772984) or vehicle (DMSO) prior to immunoblotting for markers of ERK and mTORC1 activation. (pooled protein from n=2 wells/lane) B) Small airway epithelial cells (SAECs) were subjected to volutrauma (24% biaxial stretch, 0.2 Hz, 30 min) in the presence of increasing doses of the Akt inhibitor (MK-2206) or vehicle (DMSO) prior to immunoblotting for markers of Akt and mTORC1 activation. (pooled protein from n=2 wells/lane) C-D) HBEs were stretched in the presence of CellRox Green and fluorescence was quantitated in each high powered field. Data log normally distributed, analyzed by on log_2_ transformed data. (n=18 high power fields/group) E-F) SAECs were subjected to volutrauma for 30 minutes in the presence of mitoSOX (red) and calcein AM (green) prior to quantitating the intensity of mitoSOX Red staining. Data not normally distributed, analyzed by Mann-Whitney test. (n=27 images per condition) G) Small airway epithelial cells were treated with 500 µM hydrogen peroxide (H_2_O_2_) for increasing amounts of time prior to immunoblotting for markers or ERK and mTORC1 activation. (n=1 well/lane) H) HBE cells were treated with increasing doses of glutathione ethyl ester (GSH-EE) 30 min prior to volutrauma and immunoblotting for markers of mTORC1 activation. (n=1 well/lane) *p<0.05 for all panels

### Rapamycin mitigates physiologic lung dysfunction during VILI

To determine the therapeutic potential of targeting mTORC1 activation to prevent VILI, we treated wild-type mice with the mTOR inhibitor rapamycin (5 mg/kg IP) or vehicle control at the time mechanical ventilation was initiated. Mice were subjected to simultaneous volutrauma and atelectrauma (12 cc/kg, PEEP 0 cm H_2_O) for 4 hours. Treatment with a single dose of rapamycin dramatically decreased P-S6 levels in whole lung tissue following VILI **(Figure 6A)**. Following injurious ventilation vehicle treated mice had increased lung elastance (i.e. stiffness) compared to rapamycin treated mice (**Figure 6B)**. Over the period of ventilation, rapamycin treated mice had significant lung recruitment as evidenced by an increase in inspiratory capacity that was not seen in vehicle treated mice **(Figure 6C)**. Although rapamycin prevented physiologic lung dysfunction following VILI, it did not affect recruitment of inflammatory cells **(Figures 6D-E)** or alter BAL IL-6 or KC levels. **(Figure 6F-G)**. BAL protein levels were also not different between rapamycin and vehicle treated mice **(Figure 6G)** mirroring our findings in mice with airway epithelial *Tsc2* deletion. These data demonstrate that rapamycin administered at the time mechanical ventilation is initiated decreases mTORC1 activation and prevents physiologic lung injury during mechanical ventilation independent of lung inflammation or barrier permeability.

**Figure 6.**
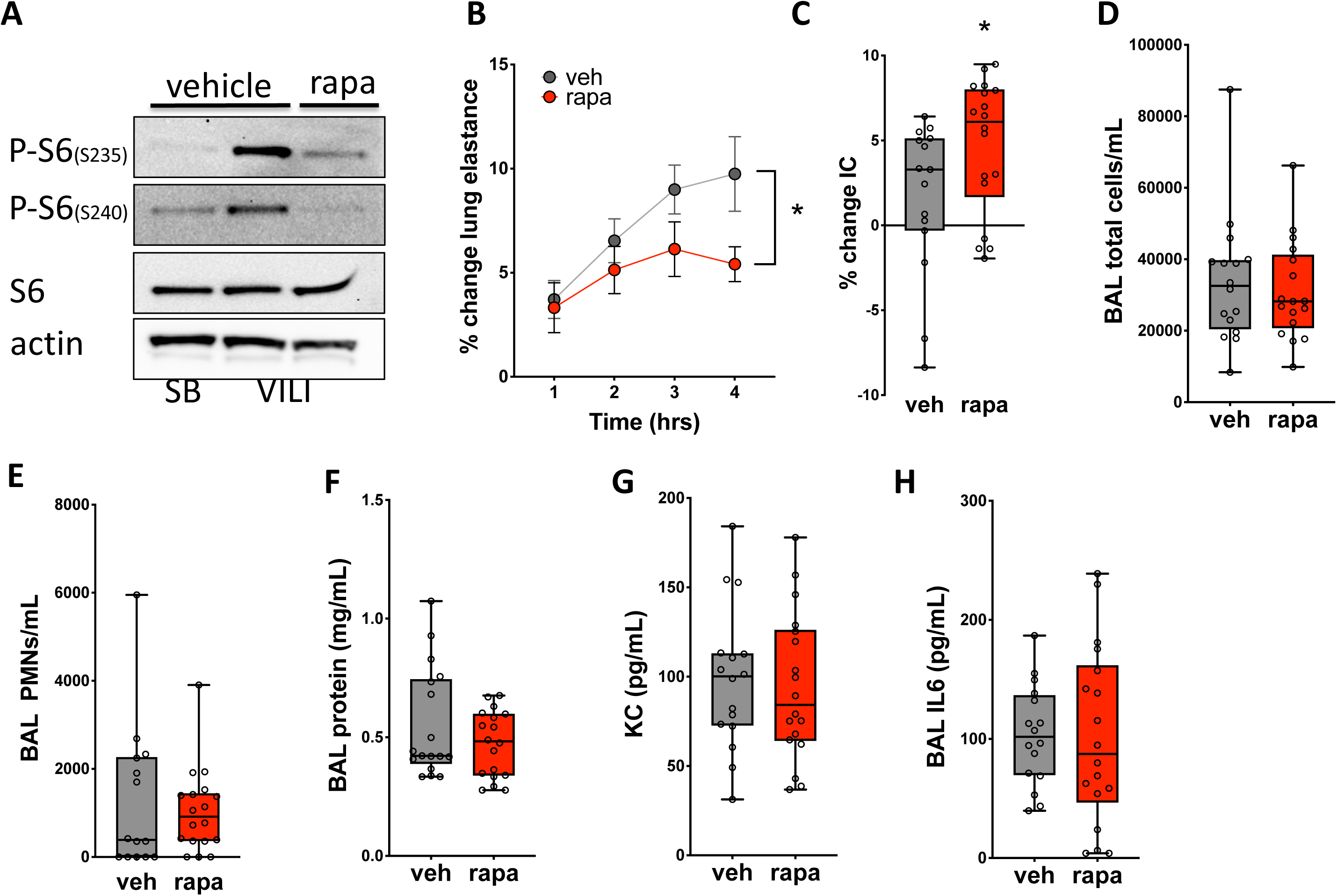
Pharmacologic mTORC1 inhibition attenuates VILI. Wild type mice were treated with rapamycin (rapa) or vehicle (veh) immediately prior to injurious mechanical ventilation (VILI, tidal volume 12 cc/kg, PEEP 0 cm H_2_O) for 4 hrs. A) Following mechanical ventilation, protein was isolated from lung tissue of ventilated (n=6) and spontaneously breathing (SB) control (n=4) mice and immunoblotted for phosphorylated S6 (P-S6). B) The change in lung elastance was measured during the 4 hr period of mechanical ventilation. *indicates that rapamycin is a statistically significant factor, p<0.05, by 2-way ANOVA, data presented as mean ± SEM. C) The percent change in inspiratory capacity (IC) was quantitated following 4 hrs of VILI. Data not normally distributed, analyzed by Mann-Whitney test, *p<0.05. Following mechanical, a bronchoalveolar lavage (BAL) was performed and total inflammatory cells (D), neutrophils (E, PMNs, n=14 vehicle, n=18 rapa), and protein levels (F) were measured. KC (G) and IL-6 (H) levels were measured in BAL fluid by ELISA. n=16 for vehicle and n=18 for rapamycin unless otherwise noted.

### mTORC1 activation exacerbates surfactant dysfunction during injurious ventilation

Since our data demonstrate that mTOR inhibition **(Figure 6)** and *Tsc2* deletion **(Figure 2)** regulate physiologic lung function without altering barrier permeability or inflammation, we hypothesized that mTORC1 activation exacerbates lung injury during VILI by impairing surfactant function. Surfactant is a complex mixture of proteins and phospholipids that decreases surface tension and prevents lung collapse under physiologic conditions. VILI and ARDS cause surfactant dysfunction and impair physiologic lung function by increasing alveolar collapse and decreasing lung compliance.(37, 38) To measure the effects of mTOR inhibition on surfactant composition we separated mouse lung surfactant into the surface-active large aggregate (LA) and inactive small aggregate (SA) fractions and measured phospholipid levels. Mice treated with rapamycin had slightly lower amounts of total and LA phospholipid. **(Figure 7A-B)** There was no difference in SA phospholipid levels or the ratio of LA/SA phospholipid **(Figure 7C-D)**. We then used a constrained drop surfactometer (CDS) (39, 40) to examine the effects of mTORC1 activation on surfactant function. This device allows for the measurement of surface tension dynamics in small volume murine samples and we validated the CDS by analyzing surface tension versus area plots in serial dilutions of recombinant surfactant (Infrasurf). As shown in **Figure 7E**, undiluted Infrasurf exhibited a large hysteresis area and near zero minimum surface tension, while diluted Infrasurf exhibited decreased hysteresis area and increased minimum surface tension. (41) After equalizing phospholipid concentrations in LA fractions, rapamycin treated mice had lower minimum surface tension compared to vehicle treated mice **(Figures 7F-H)**. These data demonstrate an association between mTORC1 activation during VILI and altered surfactant function.

**Figure 7.**
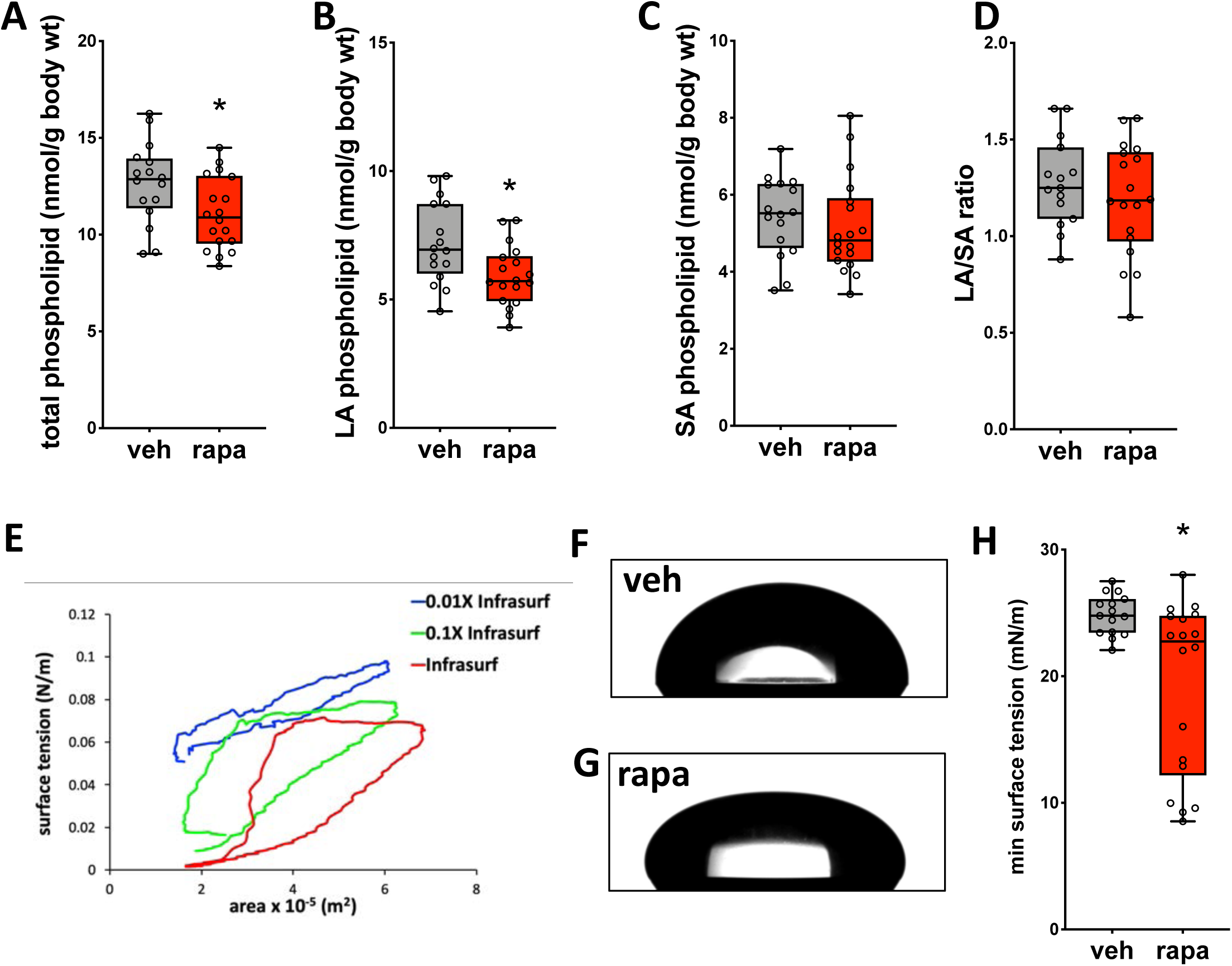
mTORC1 activation exacerbates surfactant dysfunction during injurious ventilation. A) Total phospholipid levels were measured in vehicle (veh) or rapamycin (rapa, 5mg/kg) treated mice following 4 hours of injurious ventilation (tidal volume 12 cc/kg, PEEP 0 cm H_2_O). Data normally distributed, analyzed by student’s t-test. Phospholipid levels were also measured in large aggregate (LA) fractions (B), small aggregate (SA) fractions (C), and the ratio or LA/SA phospholipid was calculated (D). Data normally distributed, analyzed by student’s t-test. E) Surface tension vs surface area plots of serial dilutions of Infrasurf measured by surfactometry. F-G) Images of minimum surface tension from LA fractions of mice treated with rapamycin or vehicle prior to VILI. H) Surface tension was measured using a constrained drop surfactometer followed by axiosymmetric drop shape analysis in LA fractions from vehicle and rapamycin treated mice. Data not normally distributed, analyzed by Mann-Whitney test. *p<0.05, n=16 for vehicle and n=18 for rapamycin.

### mTORC1 activation impairs release of the surfactant secretagogue extracellular ATP in response to volutrauma

During physiologic respiration, stretch of type I alveolar epithelial cells (AT1) stimulates surfactant release from adjacent type II alveolar epithelial cells (AT2) through the controlled release of ATP into the extracellular space and subsequent binding to purinergic receptors on AT2 cells. (42, 43) Stretch induced release of surfactant plays a key role in surfactant secretion under homeostatic conditions and is an important compensatory response to mitigate lung dysfunction during injurious ventilation. (37, 44, 45) We hypothesized that airway epithelial cells also release ATP in response to VILI but that mTORC1 activation in these cells impairs this compensatory response and exacerbates to surfactant dysfunction. To test this hypothesis we subjected SAECs and HBEs to in vitro volutrauma and found a 2-5 fold increase in the concentration of extracellular ATP released into the culture medium **(Figure 8A-B)**. A similar increase in extracellular ATP was seen when SAECs were subjected to in vitro atelectrauma **(Figure 8C)**. Interestingly, in vitro barotrauma did not increase extracellular ATP levels in either SAECs or HBEs suggesting that different mechanical forces may have unique effects on the release of extracellular ATP **(Supplemental Figure 9)**. mTORC1 inhibition with rapamycin or Torin 2 significantly increased the release of extracellular ATP following volutrauma **(Figures 8D-E)**. Given that volutrauma induced mTORC1 activation is ERK dependent, we treated HBE cells with an ERK inhibitor prior to stretch and found that ERK inhibition prior to volutrauma also increased the release of extracellular ATP **(Figure 8F)**. These data indicate that airway epithelial cells, release ATP into the alveolar space following volutrauma and atelectrauma. However, mTORC1 activation in these cells limits the release of extracellular ATP which can be enhanced by treatment with mTORC1 or ERK inhibitors. Paracrine release of ATP from airway epithelial cells may be a compensatory mechanism to increase surfactant release from adjacent AT2 cells during injurious ventilation.

**Figure 8.**
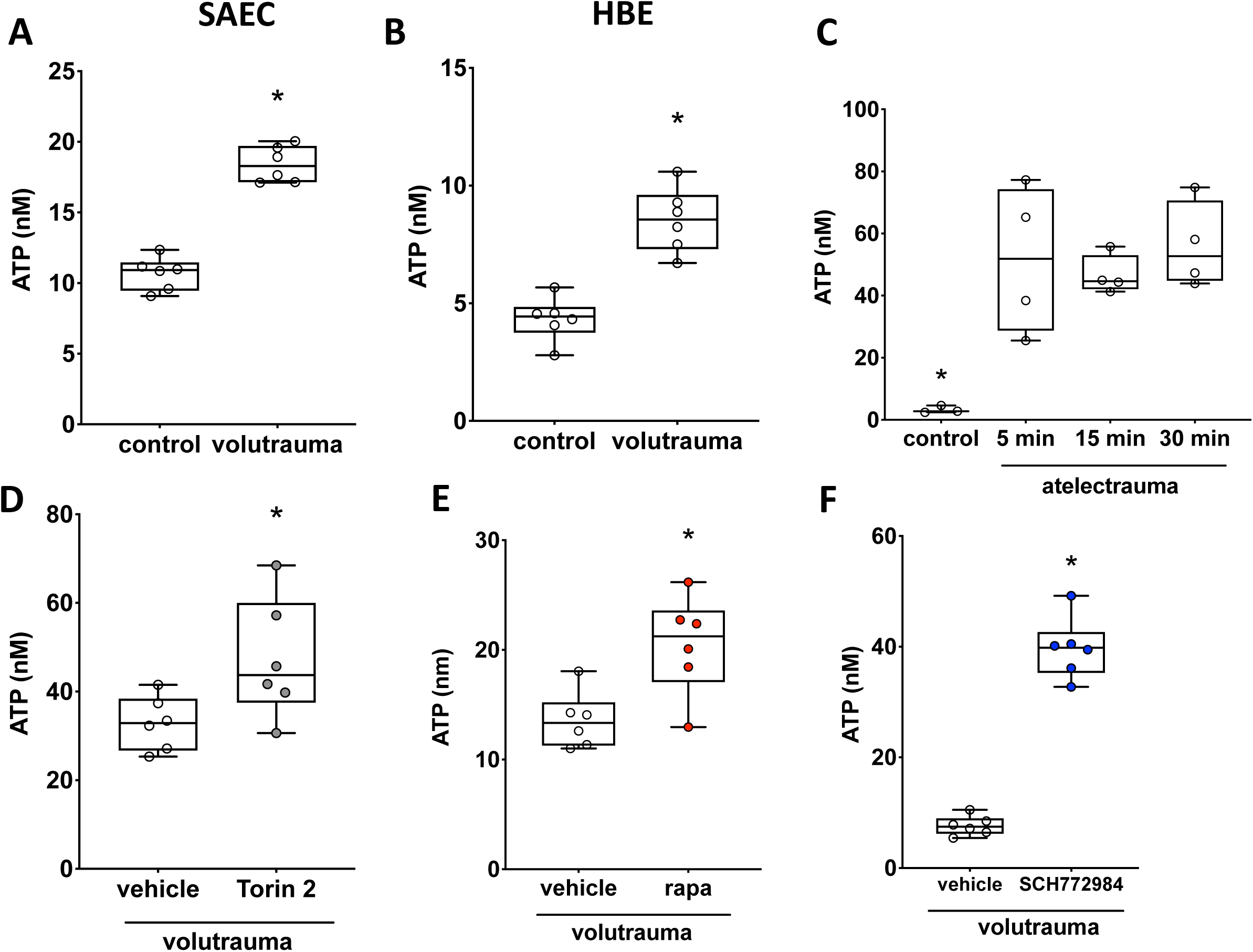
mTORC1 activation impairs release of the surfactant secretagogue extracellular ATP in response to volutrauma. A) Small airway epithelial cells (SAEC) were subjected to volutrauma (24% stretch, 0.2 Hz, 30 min) or static culture (control) prior to measuring extracellular ATP. Data normally distributed, analyzed by student’s t-test, n=6 wells/condition. B) Human bronchial epithelial cells (HBE) were subjected to volutrauma (24% stretch, 0.2 Hz, 30 min) or static culture (control) prior to measuring extracellular ATP. Data normally distributed, analyzed by student’s t-test, n=6 wells/condition. C) SAECs on polyacrylamide gels were subjected to atelectrauma or control for varying amounts of time prior to measuring extracellular ATP. Data normally distributed, analyzed by 1-way ANOVA with Sidak’s post hoc test, n=4 for all timepoints except control n=3. D-F) HBEs were treated with Torin 2 (D, 10 nM), rapamycin (E, 10 nM), SCH772984 (F, 10 µM) or appropriate vehicle control 1 hr prior to in vitro volutrauma. ATP concentration was measured in media following 30 minutes of injurious stretch. Data normally distributed, analyzed by student’s t-test, n=6 wells/group. *p<0.05 vs all groups

## Discussion

Classically, favorable growth conditions (*e.g.* presence of nutrients and growth factors) activate mTORC1, and cellular stress (*e.g.* starvation) inhibits mTORC1. (46, 47) In contrast to canonical mTORC1 activation in the setting of favorable growth conditions, we found that this pathway was activated in the lung epithelial cells during injurious mechanical ventilation in preclinical animal models, in in vitro models of VILI, and patients with ARDS. Deletion of *Tsc2*, a negative regulator of mTORC1, in airway epithelial cells increased mTORC1 activation and physiologic lung dysfunction following injurious mechanical ventilation **(Figure 2)**. Conversely, treatment with an mTOR inhibitor (i.e. rapamycin) at the time that mechanical ventilation was initiated prevented physiologic lung dysfunction **(Figure 6)**. Pharmacologic intervention at the time when MV begins is an appealing and practical therapeutic strategy that could be translated to patients with respiratory failure that require ventilatory support.

Despite changes in lung function and oxygenation in our in vivo experiments, neither genetic deletion of *Tsc2* nor pharmacologic mTOR inhibition altered lung inflammation or barrier permeability **(Figures 2 and 6)**. These findings suggested that mTORC1 activation may impair physiologic lung function during VILI by regulating surfactant function. Surfactant dysfunction is a pathophysiologic hallmark of ARDS that exacerbates lung injury in a feed forward manner, although the molecular mechanisms driving this dysfunction have not been elucidated.(48) We found that mice treated with rapamycin had improved surfactant function **(Figure 7)** that resulted in improved lung compliance **(Figure 6)**. Studies from patients with ARDS have shown decreased total phospholipid levels as well as alterations in the biochemical composition of the lipids that compose surfactant can contribute to surfactant dysfunction. (49) In addition, surfactant can be inactivated in the alveolar spaces by secreted phospholipases.(50) Interestingly, increased mTORC1 activation is associated with altered cellular lipid metabolism (51) and prolonged treatment with the allosteric mTOR inhibitor, temosirolimus, increases levels of surfactant lipid levels in mice.(52) mTOR activation has been shown to play a role in the development of neonatal lung injury which is characterized inadequate surfactant production.(53) In a mouse model of neonatal respiratory distress syndrome from hyperoxia, rapamycin treatment decreased lung injury.(54) Our data demonstrate that rapamycin treatment at the time mechanical ventilation was initiated mitigated VILI induced surfactant dysfunction **(Figure 7)**. This occurred despite rapamycin treated mice having decreased phospholipid levels in the surface active large aggregate fraction. This suggests that the mechanism by which mTOR inhibition mitigates surfactant may be by altering the phospholipid composition or other physicochemical properties of surfactant. Our data indicate that, in the short-term, mTORC1 inhibition may be a useful and feasible therapeutic strategy to mitigate surfactant dysfunction during VILI to prevent further injury.

Activation of mTORC1 has been reported in mouse models of inflammatory lung injury including endotoxin induced lung injury (15, 55, 56), hyperoxia (54), and bacterial infection (57). Similar to our findings, mTORC1 activation was also increased in alveolar and airway epithelial cells following LPS-induced lung injury. (55) In genetic mouse models in which mTOR was deleted from alveolar and airway epithelial cells, cytokine levels were decreased and pulmonary edema was increased following LPS-induced lung injury.(55) In contrast with these prior reports, we did not see differences in lung inflammation or barrier permeability with mTOR inhibition in our model of injurious ventilation. **(Figures 6)**. This may be due to inherent differences in the mechanism of lung injury, the timepoints studied, or a combination of these factors. mTORC1 activation has also been examined in inflammatory cells such as neutrophils (15) and Th17 cells (58) following endotoxin induced lung injury. Our data suggest that mTOR inhibition can mitigate lung injury during mechanical ventilation independent of inflammation.

mTOR activation has been shown to play a key role in other forms of lung injury. Pulmonary fibrosis is a condition characterized by the deposition of a stiff extracellular matrix and mechanical cues from the matrix are known to drive the pathogenesis of lung fibrosis.(59) Interestingly, expression of mTOR is increased in epithelial cells of patients with pulmonary fibrosis (60) and Chung et al. showed that rapamycin decreased mortality and fibrosis in a murine model of radiation induced fibrosis.(61) In contrast to the injurious role of mTORC1 activation in LPS and fibrosis induced lung injury, mTORC1 activation appears to play a protective role in the development of cigarette smoke induced emphysema. Yoshida et al. showed that a stress response protein, RTP801, is induced by cigarette smoke and mediates the development of emphysema, in part, by inhibiting mTORC1. (14) More recently, Houssaini et al. showed that activation of mTORC1 and mTORC2 increased senescence in lung cells from patients with COPD or in mice with hyperactivated mTOR in alveolar epithelial cells.(62) These data demonstrate the complex and pleiotropic role of mTOR activation during different mechanisms of lung injury and suggest that a deeper understanding of the pathobiologic mechanisms that lead to lung injury will be important for developing targeted therapies for these patients.

Activation of mTORC1 by mechanical stimuli has been described in skeletal and cardiac muscle in the context of compensatory hypertrophy,(11–13, 63) and as a key regulator of bone and cartilage growth during development. (64, 65) Canonical activation of mTORC1 by growth factors involves phosphorylation and subsequent inhibition of tuberin, a constitutive inhibitor of mTORC1 encoded by the *Tsc2* gene, by upstream kinases such as ERK and Akt.(27) Prior data regarding the role of Akt in mechanical mTORC1 activation in skeletal muscle have been mixed. Bodine et al. used in vivo rat models to demonstrate that Akt was activated following contractile stress in skeletal muscle in a model of compensatory hypertrophy and inhibited in a model of muscle atrophy. (13) Using an ex vivo model of skeletal muscle loading, Hornberger et al. showed that tensile stretch activated mTORC1 in an Akt-independent fashion.(66) Similarly, our data demonstrate that mechanical activation of mTORC1 in lung epithelial also occurs in a Akt-independent manner **(Figure 5)**.

In contrast to canonical activation of mTORC1, biomechanical activation in muscle can occur in the absence of growth factors (66) via activation of the ERK pathway. (12) Similar to force induced mTORC1 activation in muscle, activation of the complex in lung epithelial cells did not require the presence of exogenous growth factors **(Figures 4-5)**. Despite being growth factor independent, our data demonstrate that injurious stretch in lung epithelial cells rapidly activates the ERK1/2 pathway and that volutrauma induced mTORC1 activation is ERK-dependent **(Figure 5)**. A variety of second messengers can activate ERK1/2 in lung epithelial cells. Given that physical forces during MV are known to release reactive oxygen species (67, 68) and mTORC1 can be regulated in a redox sensitive fashion (69, 70) we examined the role of ROS in activating mTORC1 during VILI. Our data demonstrate that exogenous ROS (i.e. H_2_O_2_) or volutrauma induced release of cellular ROS rapidly activate mTORC1 **(Figure 5, Supplemental Figure 7)** and that this activation is inhibited by the antioxidant GSH **(Figure 5)**. This finding is consistent with prior data showing that short term exposure to ROS can activate mTORC1 in a variety of cells lines (33, 34) and demonstrates this effect for the first time in primary human lung epithelial cells. Excessive production of free radicals in the context of lung injury has been shown to mediate the development of inflammation, barrier permeability, and cell death.(67, 71) Our data uncover a novel mechanism by which ROS drive the development of lung injury by acting as second messengers that activates mTORC1 signaling.

Several mTOR inhibitors are currently FDA approved for the prevention of rejection after organ transplant, for the treatment of tuberous sclerosis complex, and to prevent lung function decline in patients with lymphangioleiomyomatosis.(72) In our in vivo studies we administered rapamycin immediately prior to the initiation of mechanical ventilation which is particularly relevant for preventing lung injury in the context of mechanical ventilation. In summary, our data demonstrate that injurious physical forces during mechanical ventilation activate mTORC1 in lung epithelial cells and that activation of this pathway in this context drives the development of lung injury. Future studies will be needed to further explore the role of mTORC1 inhibition to prevent lung injury in ICU patients undergoing mechanical ventilation.

## Methods

### Reagents

Rapamycin and Torin 2 were purchased from LC Laboratories. SCH772984 and MK-2206 were purchased from SelleckChem. ARL 67156 was purchased from Tocris. Glutathione ethyl ester (buffered to pH 7.4 prior to use), antimycin A, perchloric acid, and ammonium molybdate were purchased from Sigma-Aldrich. Ascorbic acid was purchased from VWR. MitoSOX, CellROX Green, calcein, and propidium iodide were purchased from Invitrogen. Calfactant (Infasurf) was purchased from The Ohio State University Wexner Medical Center pharmacy. A complete list of antibodies can be found in the Supplemental Materials.

### Cell culture

Primary human small airway epithelial cells were obtained from Lonza (Walkersville, MD) and PromoCell (Heidelberg, Germany) and grown in the manufacturer’s suggested growth medium. Primary human bronchial epithelial cells were also obtained from Lonza and grown in the recommended growth medium. About 12 hours prior to in vitro models of VILI, cells were washed with warm PBS and changed to the corresponding basal medium with penicillin/streptomycin (100 U/mL, 100 μg/mL; Gibco) and amphotericin B (0.25 μg/mL, Gibco) but without growth factors or serum.

### Mouse tracheobronchial epithelial cell isolation

Tracheobronchial epithelial cells were isolated from *CC10*^Cre^/*Tsc2*^flox/flox^ and *Tsc2*^flox/flox^ mice as previously described.(22) Briefly, the trachea and proximal bronchi were isolated and digested in 0.15% pronase (Roche) overnight at 4 °C. The digestion was stopped with 10% FBS (Gibco) and cells were pelleted for DNase treatment (Sigma-Aldrich). Following adherence purification, cells in the suspension were collected for analysis by immunoblotting.

### Mouse strains

Mice expressing homozygously floxed *Tsc2* alleles (*Tsc2*^flox/flox^) and mice expressing Cre recombinase under the control of the CC10 promoter (*CC10*^Cre^) were generously provided by Dr. Lisa Henske. *CC10*^Cre^ mice were bred with *Tsc2*^flox/flox^ to generate mice with airway epithelial specific deletion of Tsc2. *mT/mG* reporter mice were obtained from Jackson Laboratory (#007576). *CC10*^Cre^ mice were bred to *mT-mG* reporter mice(21) to confirm airway epithelium-specific expression of the Cre recombinase. Genotyping was performed by Transnetyx Inc. using real-time PCR for both *Tsc2* and Cre alleles. All mice were backcrossed to a pure C57BL/6 background which was confirmed by genome scanning performed by Jackson Laboratory. Wild-type C57BL/6 mice (#000664) were obtained from Jackson Laboratories (Bar Harbor, ME). Deletion of *Tsc2* was confirmed by immunostaining lung sections and for phosphorylated ribosomal S6 protein and immunoblotting for tuberin with protein lysate from isolated tracheobronchial epithelial cells. All mice were housed in pathogen free conditions in vivariums at Brigham and Women’s Hospital and The Davis Heart and Lung Research Institute. Mice were used for experiments at 6-12 weeks of age and provided food and water ad lib.

### Murine models of ventilator induced lung injury

Mice were sedated with a combination of ketamine (Vedco, St. Joseph, MS) and xylazine (Akorn, Lake Forest, IL) or pentobarbital (Akorn, Lake Forst, IL). For experiments using rapamycin, this was administered by intraperitoneal injection following induction of anesthesia. A tracheostomy tube was place and mice were placed on a warming pad to avoid hypothermia and were mechanically ventilated for varying periods of time using a flexiVent (SCIREQ, Montreal, Canada). Oxygen saturations were measured by pulse oximetry using the MouseOx system (Starr Life Sciences, Oakmont, PA) and EKG tracings were monitored using the flexiVent system. Lung physiology parameters were measured at baseline and hourly thereafter for the duration of mechanical ventilation. Standardized recruitment maneuvers were performed prior to each physiologic measurement to prevent atelectasis and standardize volume history.(73) For experiments using the CLP/VILI model, mice were anesthetized and underwent cecal ligation and puncture (23 gauge needle, 1 hole, 50% ligation) or sham laparotomy as described previously(74) 24 hours prior to mechanical ventilation. At the end of the mechanical ventilation protocol, animals were sacrificed by anesthetic overdose and a bronchoalveolar lavage (BAL) was performed by instilling 1 mL of sterile PBS twice. The BAL fluid was then centrifuged for 10 min at 400 *g* at 4°C. Pelleted cells were treated with RBC lysis buffer (Alfa Aesar), counted by an automated cell counter (Bio-Rad), and cytospins were performed following by Diff-Quick staining (Fisher Scientific) to obtain differential cell counts.

### Surfactant phospholipid and function measurements

Following mechanical ventilation, lungs were lavaged with 1 mL of 0.9% NaCl solution 3 times for a total volume of 3 mL. The BAL fluid was then centrifuged for 10 min at 400 *g* at 4°C. The supernatant was centrifuged at 40000 *g* for 18 min to obtain the small aggregate fraction (supernatant) and the large aggregate fraction (pellet), which was concentrated by resuspension in a small volume of 0.9% NaCl solution. Phospholipids were extracted from fractionated pulmonary surfactant samples by the Bligh and Dyer method.(75) Phosphorus content was measured by colorimetry following perchloric acid digestion and treatment with ammonium molybdate and ascorbic acid.(76) Surfactant surface tension was measured using a constrained drop surfactometer (CDS) based on a prior design (39) that consisted of a modified goniometer, digital camera (ramé-hart, Succasunna, NJ) and custom-built pedestal. The CDS was manufactured with 316 stainless steel and consists of a circular pedestal with a sharp 60° edge to prevent film leakage. 35 μL droplets of resuspended large aggregate fractions were placed on the pedestal of the CDS. The droplet was oscillated using a programmable syringe pump at a rate of 6 seconds per cycle and compressed to ∼50% of the initial surface area. This was repeated for a total of 15 cycles per large aggregate sample. A custom-written MATLAB code was used to analyze the droplet shape in each frame to obtain surface tension and surface area measurements. First, the Sobel method was used to detect the droplet edge in each frame. Next, the differential equations governing force balances for an arbitrary axisymmetric droplet shape (77) were integrated to determine a theoretical droplet shape based on the surface tension, tip radius of curvature and angle of inclination parameters. Finally, a non-linear least-squared regression algorithm was used to vary the surface tension and tip radius of curvature parameters until the theoretical droplet shape matched the measured droplet shape. The droplet shape was also integrated to determine the surface area in each frame. This algorithm results in surface tension vs time and surface tension vs area hysteresis loops that were further analyzed for minimum surface tension.

### In vitro volutrauma model

Cells were cultured on pronectin-coated BioFlex culture plates (FlexCell International Corporation) and grown to confluence. Cells were subjected to varying degrees of biaxial cyclic stretch delivered via a Flexcell Strain Unit FX-5000 (FlexCell) for varying amounts of time prior to protein isolation or imaging. Control cells were placed in the stretching device, but not subjected to mechanical stretch.

### In vitro atelectrauma model

An in vitro model of atelectrauma was used as previously described.(28, 78) Briefly, polyacrylamide gels containing 10% w/w acrylamide (Bio-Rad) and 0.25% w/w bis-acrylamide (Bio-Rad) were fabricated on 40 mm glass slides (20 kPa, 200 µm thickness). Polyacrylamide gels or glass slides were coated with collagen type I (Sigma-Aldrich) at a concentration of 5 μg/cm^2^ following surface activation of the glass slide with sulfo-SANPAH (Thermo Scientific). Cells were cultured on the gels or glass slides and placed in a microfluidic chamber (Bioptechs Inc.) with a rectangular silicone gasket to create the channel for cyclic propagation of media at a linear velocity of 30 mm/s for varying amounts of time.

### In vitro barotrauma model

Oscillatory pressure was applied as described previously. (79) Briefly, cells were cultured on Transwell inserts (0.4 µm pore size, PET; Corning Inc.) and grown to confluence. Media was removed from the apical chamber of the Transwell and a rubber stopper was inserted into the culture well in order to hermetically seal the apical chamber. Tubing was threaded through the rubber stopper and connected to a manometer and small animal ventilator (Harvard Apparatus). The ventilator was set to 0.2 Hz at a magnitude of 20 cm of H_2_O of oscillatory pressure.

### Extracellular ATP measurements

Cells were subjected to in vitro models of VILI in the presence of an ectonucleotidase inhibitor ARL 67156 (50 µM) to prevent ATP degradation. Media was collected following in vitro models of VILI and extracellular ATP was measured using a luciferase-luciferin assay (Invitrogen).

### Measurements of cellular reactive oxygen species

Cells were grown to confluence and changed to basal medium 24 hours prior to in vitro volutrauma. MitoSOX (5 µM) and calcein (4 µM) was added to each well 30 minutes prior to injury. Antimycin A (20 µM) was used as a positive control. Following stretch cells were washed 3 times with PBS, membranes were cut out of the Flexcell plates and inverted into a 60 mm dish containing a small amount of basal medium immediately prior to obtaining fluorescence images. For MitoSOX studies, nine separate 100X images were captured from each membrane. Cells were defined in ImageJ using an ROI detection algorithm on the calcein stained images and MitoSOX intensity was quantitated in ImageJ for each cell within capture. Average intensity per cell was calculated for each capture. For CellROX experiments, average fluorescence per high power field was quantitated using ImageJ.

### Immunoblotting

Cells or lung tissue were lysed in RIPA buffer (VWR) supplemented with protease (Roche Diagnostics) and phosphatase inhibitors (Sigma-Aldrich). Cells were mechanically disrupted using a rubber scraper followed by a freeze/thaw cycle. Lung tissue was disrupted using a tissue homogenizer. Protein extracts were centrifuged to pellet insoluble debris, protein concentration was measured by BCA assay (Thermo Scientific), and protein concentrations were equalized prior to boiling in the recommended sample buffers supplemented with 2.5% beta-mercaptoethanol (Bio-Rad). SDS-PAGE electrophoresis was performed using 4-12% gradient bis-tris gels or 16% tricine gels (Invitrogen) and proteins were transferred to nitrocellulose membranes (Bio-Rad). Membranes were incubated in primary antibodies overnight at 4°C following blocking in 5% w/v nonfat dry milk in TBST. After incubation with HRP-conjugated secondary antibodies, proteins were visualized using SignalFire Elite ECL Reagent (Cell Signaling Technology) and imaged using a ChemiDoc XRS+ System (Bio-Rad). Quantitative assessment was performed using Image Lab software (Bio-Rad).

### Immunohistochemistry and image analysis

Mouse lungs were perfused free of blood with 10 ml of sterile saline and inflated with 10% formalin to a pressure of 30 cm H_2_O prior to overnight fixation. Following fixation lung tissue was embedded in paraffin. For immunostaining lung sections from mice and humans were deparaffinized and rehydrated prior to boiling in antigen retrieval buffer (10 mM Na citrate/0.5% Tween 20) for 30 minutes. Tissue was permeabilized with 0.3% Triton X and quenching and blocking was performed using a cell and tissue staining kit (R&D Systems) according to the manufacturer’s protocol. Tissue was incubated with anti-phospho-S6 antibody (Ser235/236) in 2% goat serum in PBS overnight at 4°C. Non-immune serum and no primary antibody controls were performed with each experiment. H&E staining was performed using standard techniques. Blinded review was performed by a veterinary pathologist and a lung injury score was calculated.(80) Stained sections were scanned at The Ohio State Wexner Medical Center Digital Imaging Core using a NanoZoomer 2.0-HT Scan system (Hamamatsu Photonics) to generate digital whole-slide images. Quantification of the digital images was performed using the VIS software suite (Visiopharm, Hoersholm, Denmark). A single ROI was then made using a tissue detection application to measure a mean DAB stain intensity of all tissue including within each tissue section. Non-tissue areas within tissue sections such as airway and vascular lumens were excluded from the mean intensity measurement.

### BAL protein and cytokine analysis

Protein concentrations were measured in BAL fluid using a standard BCA assay (Thermo Scientific) and various analytes including IL-6, KC, and VEGF-A were measured by ELISA (R&D Systems) according to the manufacturer’s instructions.

### Study approval

All animal studies were approved by the Institutional Animal Care and Use Committee at The Ohio State University (2013A00000105-R2) and performed in accordance with NIH guidelines. Animals were randomly assigned to treatment groups and used at 6-12 weeks of age unless otherwise noted. Lung tissue was obtained from the Brigham and Women’s Department of Pathology and clinical data were collected from subjects under an approved IRB protocol 2015P002273.

### Statistics

Data were analyzed using GraphPad Prism versions 7 & 8. Data are represented as box and whisker plots (median ± min/max) unless otherwise noted. Experimental sample size was determined by performing a pilot experiment and the sample mean and standard deviation estimates were used to calculate the final sample size with power of 0.8 and α = .05. Normality and log-normality were tested using the Shapiro-Wilk test. For comparisons between 2 groups, a Student’s 2-tailed t test and Mann-Whitney test were used for parametric and non-parametric data respectively. For comparisons between multiple groups, a 1-way or 2-way ANOVA was performed depending upon the number of experimental groups. A p-value *<* 0.05 was considered statistically significant. Outliers were identified using non-linear regression via the ROUT method with a Q threshold of 1%.(81) For the clinical data, association analyses between pairs of variables were conducted with Fisher’s exact tests (for categorical variables) and two-tailed t-tests or Kruskal-Wallis tests (for continuous variables as appropriate based on the normality of the data) using SAS version 9.4 (Cary, NC).

## Author Contributions

JAE and RMB conceived and designed the study. JAE, HL, QF, AS, WZ, CI, CMB, PP, MPV, RKP, DAM, DBB, AH, HCL, NHC, SNG, JWC, RDH, and LMS acquired, analyzed, and interpreted data. JAE and HL drafted the manuscript. JAE, HL, RMB, JWC, SNG, EPH, and RDH critically revised the manuscript for intellectual content. All authors approved the final version of the manuscript and agree to be accountable for all aspects of the study.

## Acknowledgments

This work was partially supported by NIH Grants K08 GM102695 (JAE), R56 HL142767 (JAE, SNG, RDH), R01 HL142093 (RMB), and an OSU Presidential Fellowship (CMB). This work was also supported by vouchers from the Ohio State Center for Clinical and Translational Science (UL1TR002733). We would like to thank the OSU Comparative Pathology and Mouse Phenotyping Core and Dr. Sue Knoblaugh for their assistance in preparing and staining, and interpreting histologic sections, The Ohio State University College of Medicine Medical Scientist Training Program, and the Davis Heart and Lung Research Institute at The Ohio State University. The *Tsc2* flox and CC10-Cre mice were generously provided by Dr. Lisa Henske.

