## Supplemental Materials for "Mechanosensitive activation of mTORC1 mediates ventilator induced lung injury during the acute respiratory distress syndrome"

### Supplemental Methods

Supplemental Table 1. List of antibodies

| NAME | COMPANY | PRODUCT # |
|---|---|---|
| Phospho-S6 Ribosomal Protein (Ser235/236) | Cell Signaling Technology | 2211 |
| Phospho-S6 Ribosomal Protein (Ser240/244) | Cell Signaling Technology | 2215 |
| S6 Ribosomal Protein | Cell Signaling Technology | 2217 |
| Phospho-p44/42 MAPK (Erk1/2) (Thr202/Tyr204) | Cell Signaling Technology | 4370 |
| p44/42 MAPK (Erk1/2) | Cell Signaling Technology | 4695 |
| Phospho-p70 S6 Kinase (Thr389) | Cell Signaling Technology | 9205 |
| p70 S6 Kinase | Cell Signaling Technology | 9202 |
| Phospho-4E-BP1 (Thr37/46) | Cell Signaling Technology | 2855 |
| 4E-BP1 | Cell Signaling Technology | 9452 |
| Phospho-Akt (Thr308) | Cell Signaling Technology | 13038 |
| Phospho-Akt2 (Ser474) | Cell Signaling Technology | 8599 |
| Akt (pan) | Cell Signaling Technology | 4691 |
| β-Actin | Cell Signaling Technology | 3700 |
| Anti-mouse IgG, HRP-linked | Cell Signaling Technology | 7076 |
| Anti-rabbit IgG, HRP-linked | Cell Signaling Technology | 7074 |

Supplemental Data

**Supplemental Table 1. Baseline characteristics of mechanically ventilated patients from pathology tissue bank.**

| | Normal (N=5) | DAD (N=5) | P-value |
|---|---|---|---|
| Age (mean ± SD) | 56 ± 11 | 62 ± 18 | 0.6 |
| Sex (no. female, n, %) | 1 (20) | 1 (20) | 1.0 |
| Race (no. white, n, %) | 5 (100) | 4 (80) | 1.0 |
| Ever Smoker | 3 (60) | 2 (67)* | 1.0 |
| Pack Years Smoking (median, IQR) | 5 (0, 40) | 16 (0, 17)* | 1.0 |
| Heart Disease (no., %) | 0 | 3 (60) | 0.17 |
| Lung Disease (no., %) | 1 (20) | 1 (20) | 1.0 |
| CKD (no., %) | 0 | 3 (60) | 0.17 |
| Liver Disease (no., %) | 0 | 1 (20) | 1.0 |
| Malignancy (no., %) | 2 (40) | 1 (20) | 1.0 |
| Diabetes (no., %) | 0 | 3 (60) | 0.17 |
| Immune Suppression (no., %) | 0 | 1 (20) | 1.0 |

*No smoking data for 2 of the subjects

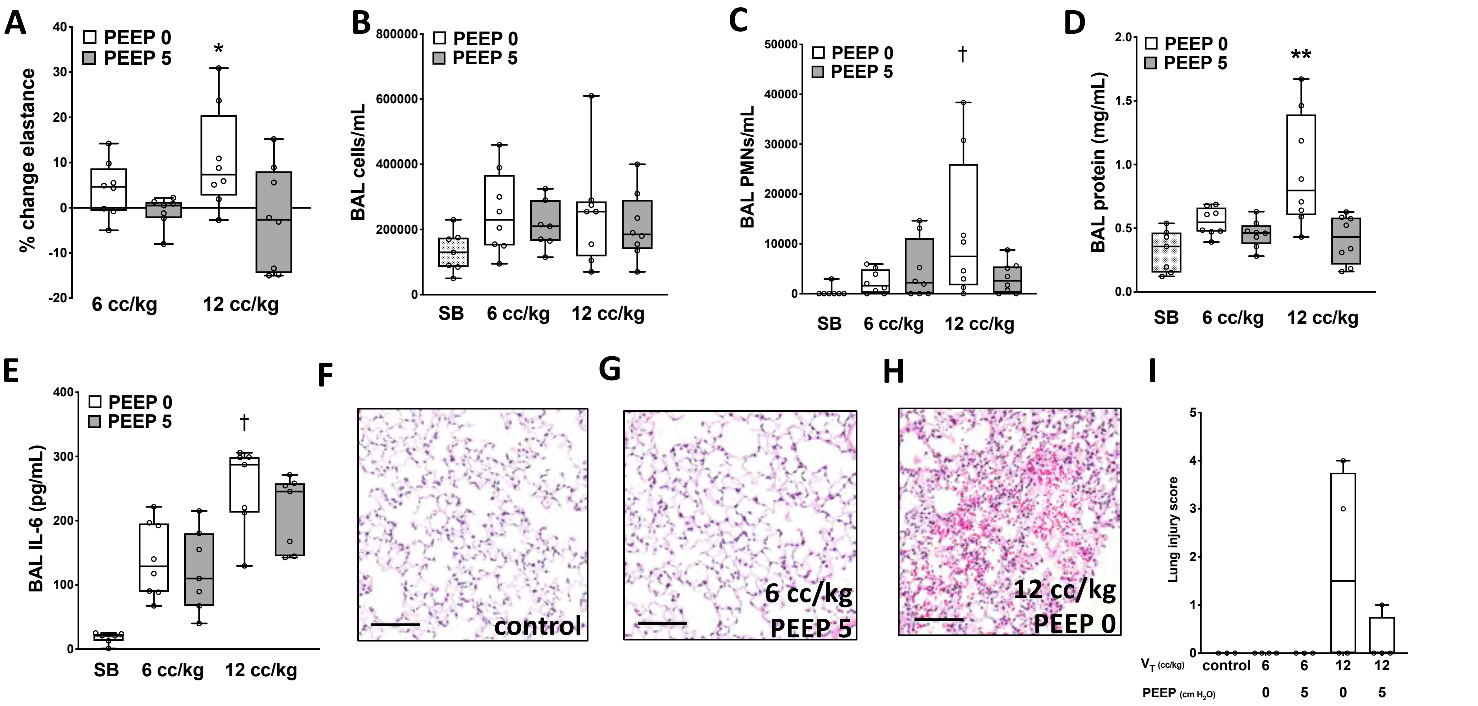


**Supplemental Figure 1. Volutrauma and atelectrauma induce lung injury in mice.** Wild type mice (n=8/group) were mechanically ventilated with high (12 cc/kg) tidal volume ($V_T$) to induce volutrauma or low $V_T$ (6 cc/kg) with or without positive end expiratory pressure (PEEP) for 4 hrs. A) Mice subjected to simultaneous volutrauma ($V_T$ 12 cc/kg) and atelectrauma (PEEP 0 cm $H_2O$) had significantly higher lung elastance (*i.e.* stiffness) compared to all other groups of mice. B-C) Total BAL cell counts were not different between groups but there were significantly more BAL neutrophils in mice ventilated with high $V_T$ without PEEP compared to spontaneously breathing (SB) control mice (n=7). Mice subjected to combined volutrauma and atelectrauma also had a significant increase in BAL protein (D) and interleukin-6 (E) levels. H&E images of lung tissue from SB control mice (F), mice ventilated with low $V_T$ (6 cc/kg) and PEEP (G), and mice subjected to simultaneous volutrauma and atelectrauma (H). I) Lung injury score was assessed by a blinded veterinary pathologist using H&E stained images (black bar=100 μm). *p<0.05 vs 12 cc/kg, PEEP 5 by 1-way ANOVA with Tukey's post-hoc test; †p<0.05 vs SB controls by 1-way ANOVA with Tukey's post-hoc test; **p<0.05 vs all groups by 1-way ANOVA with Tukey's post-hoc test.

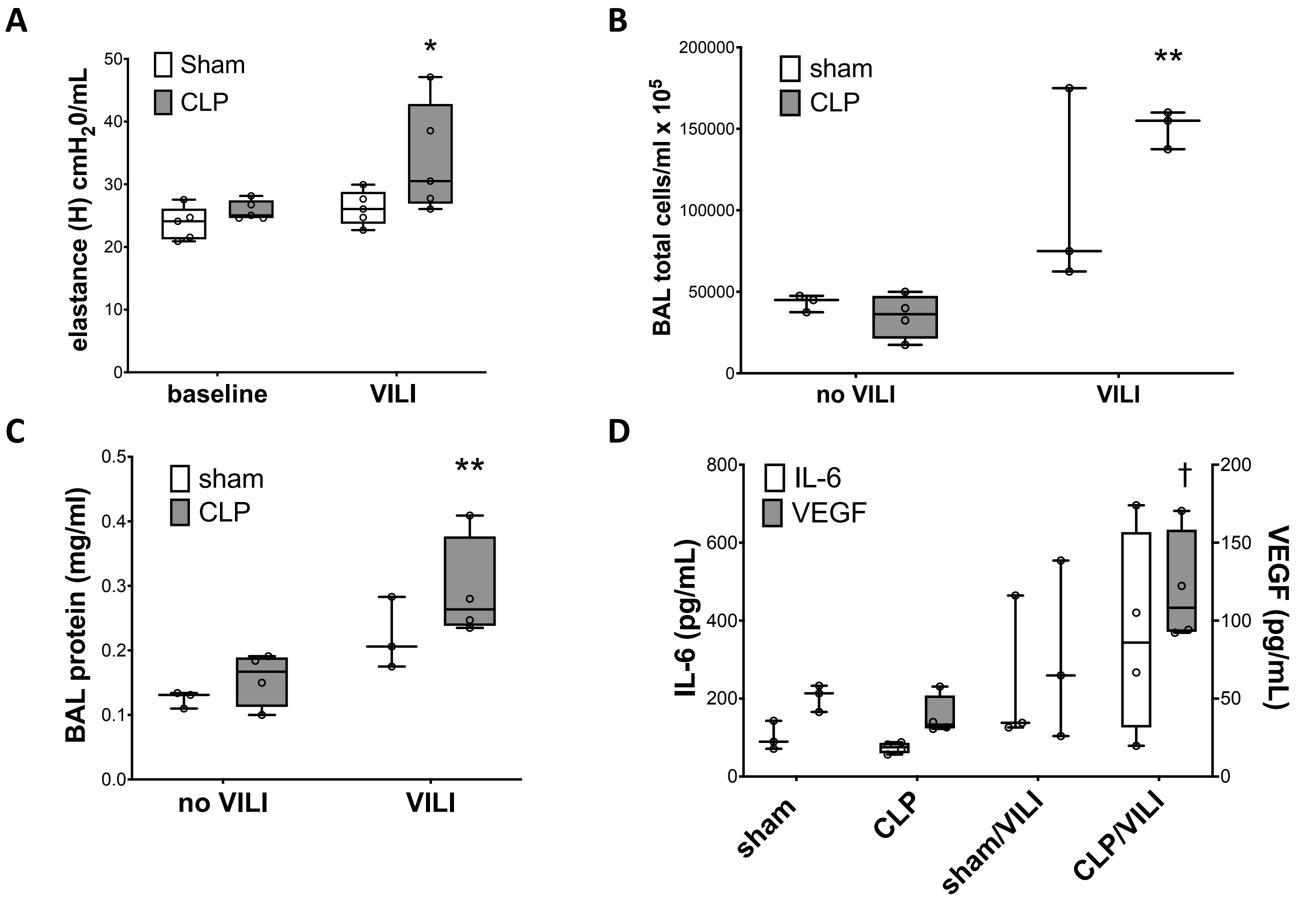


**Supplemental Figure 2. Injurious mechanical ventilation following polymicrobial sepsis exacerbates lung injury.** Wild type mice were subjected to cecal ligation and puncture (CLP, 23 gauge, 1 hole, 50% ligation) or a sham laparotomy. Mice were allowed to recover for 24 hours and were re-anesthetized and subjected to injurious mechanical ventilation (VILI-$V_T$ 12 cc/kg, PEEP 2.5 cm $H_2O$) for 4 hrs. A) Lung elastance was measured at the time mechanical ventilation was initiated (baseline) and hourly after that. CLP/VILI mice had significantly higher lung elastance at 4 hours compared to sham/VILI mice. (n=5/group, *p<0.05 vs sham/VILI by 2-way ANOVA with repeated measures and Bonferroni post-hoc test). The combination of CLP and VILI (n=4) also significantly increased the number of BAL inflammatory cells (B), BAL total protein concentration (C), and BAL VEGF levels compared to sham/VILI (n=3) mice. **p<0.05 vs sham & CLP by 2-way ANOVA with Bonferroni post hoc test. †p<0.05 vs CLP by 2-way ANOVA with Bonferroni post hoc test.

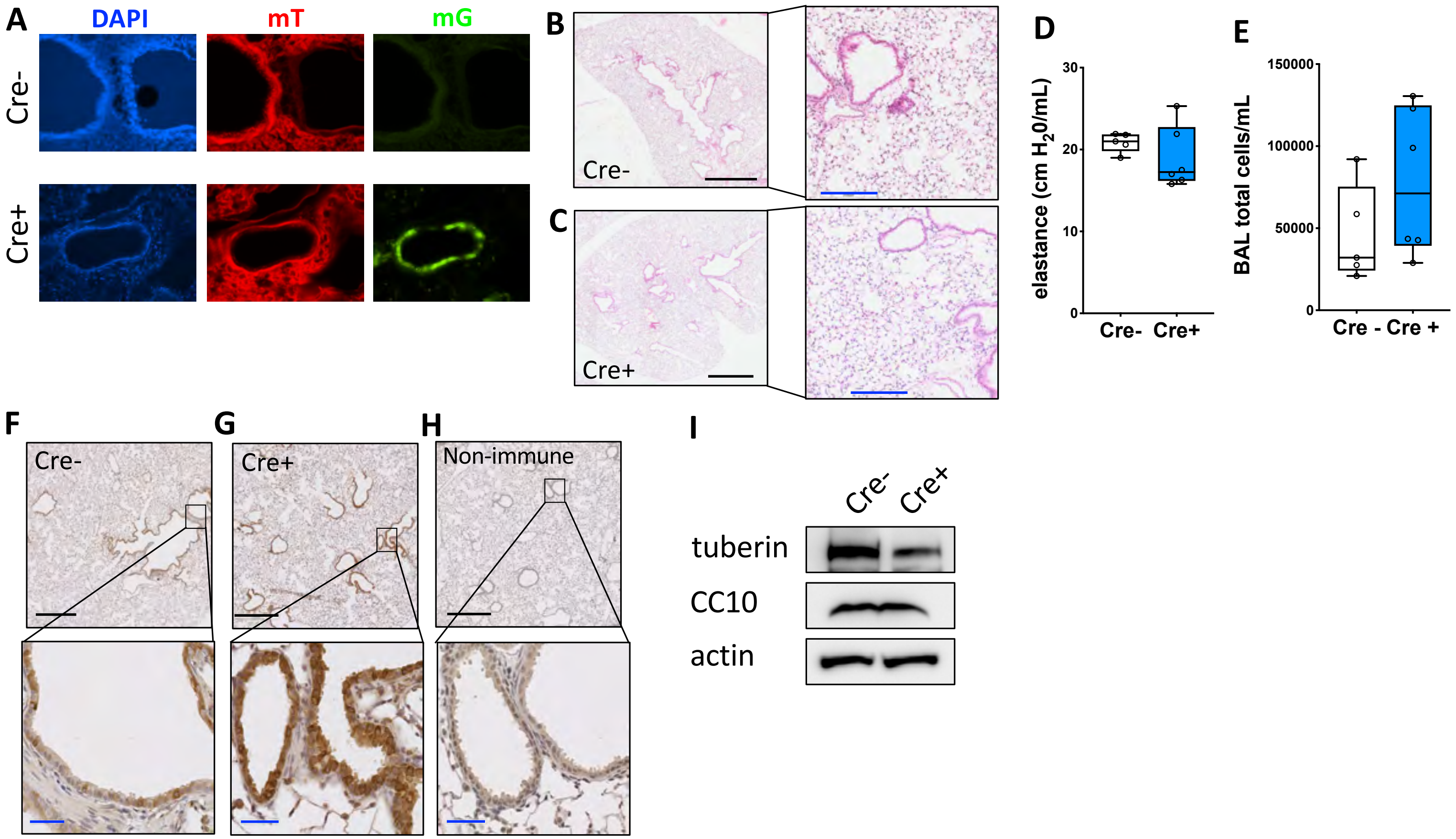


**Supplemental Figure 3. Airway epithelial *Tsc2* deletion increases mTORC1 activation but does not alter lung morphology or function in uninjured mice.** A) Immunofluorescence images of lung tissue from mT/mG transgenic mice bred to mice expressing Cre recombinase under the control of the CC10 promoter (Cre+) or control mice without Cre recombinase (Cre-). B-C) Representative photomicrographs of H&E stained lung tissue from spontaneously breathing mice with airway epithelial *Tsc2* deletion (Cre+) or controls (Cre-). (black line 1 mm, blue line 200 µm) D-E) Lung elastance (n=5 for Cre-, n=6 for Cre+) and BAL cells (n=5 for Cre-, n=7 for Cre+) from mice with airway epithelial *Tsc2* deletion (Cre+) and Cre- controls. F-G) Representative photomicrographs of lung sections from spontaneously breathing Cre- and Cre+ mice stained for phosphorylated ribosomal S6 (P-S6, Ser235/236) or non-immune rabbit serum (H, black line 500µm, blue line 50µm) I) Immunblots from isolated tracheobronchial epithelial cells from spontaneously breathing Cre- and Cre+ mice for tuberin (encoded by *Tsc2* gene), CC10, and actin. (pooled protein from n=2 mice/lane)

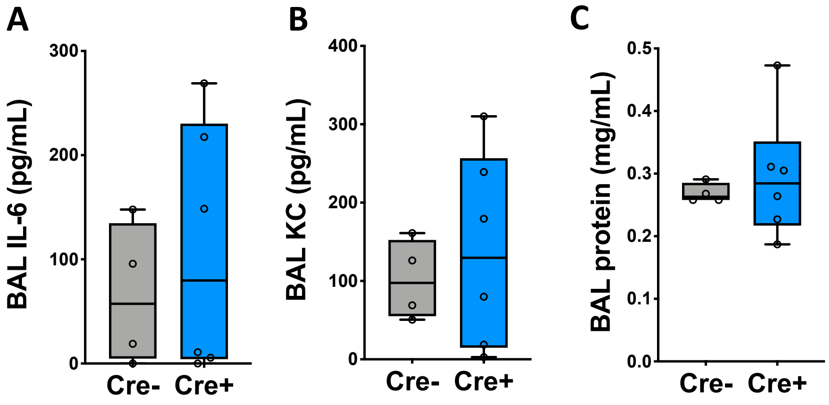


**Supplemental Figure 4. BAL IL6, KC, and protein levels are not different in spontaneously breathing mice with airway epithelial *Tsc2* deletion.** BAL IL-6 (A), KC (B), and total protein levels from spontaneously breathing Cre- and Cre+ mice. (n=4 for Cre-, n=6 for Cre+)

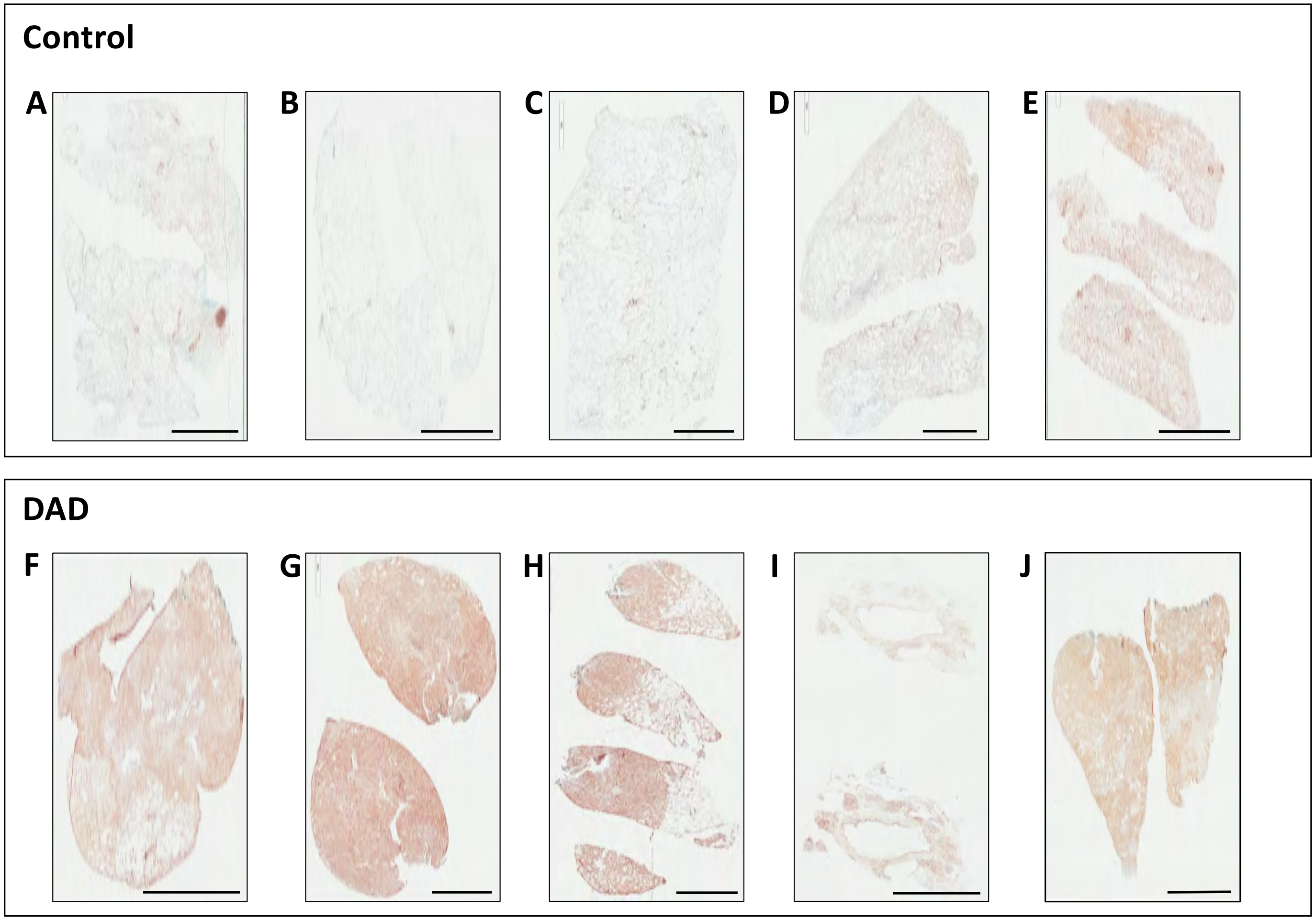


**Supplemental Figure 5. mTORC1 is activated in lung tissue from mechanically ventilated patients with diffuse alveolar damage.** Whole slide Images of P-S6 (Ser235/236) stained lung tissue from control subjects (A-E) and from patients with diffuse alveolar damage (DAD, F-J). black bar=5 mm

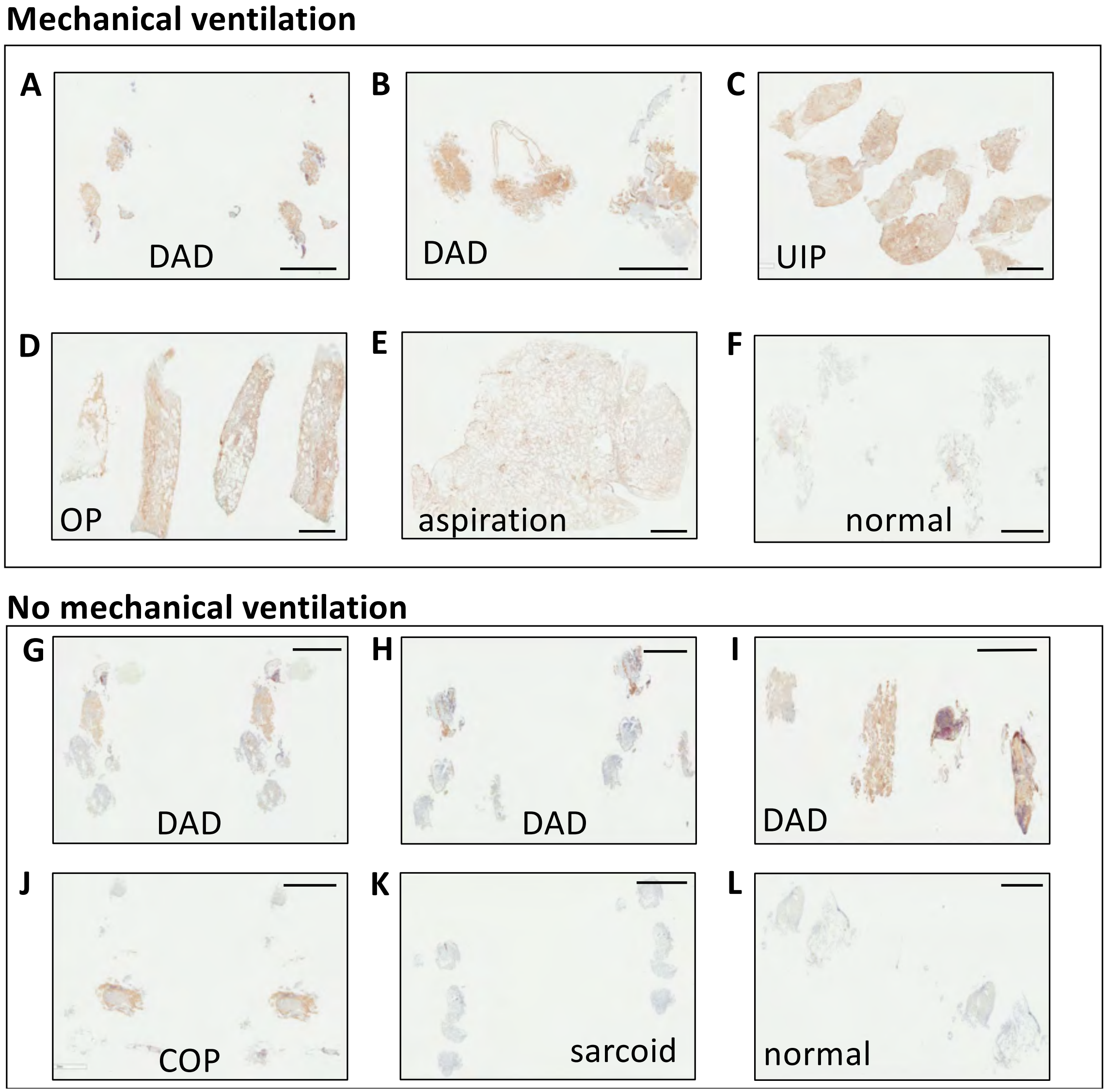


**Supplemental Figure 6. Mechanically ventilated patients have increased mTORC1 activation in lung tissue compared to spontaneously breathing patients.** Low power images of lung tissue stained for P-S6 (Ser235/236) from mechanically ventilated patients with diffuse alveolar damage (DAD) on transbronchial biopsy (A, B), other types of lung injury on surgical lung biopsy (C, UIP-usual interstitial pneumonia; D, OP-organizing pneumonia; E, aspiration pneumonitis), or no specific pathology (F). Photomicrographs of P-S6 (Ser235/236) stained lung tissue obtained by transbronchial biopsy from patients with DAD (G-I), cryptogenic organizing pneumonia (COP, J), sarcoidosis (K), or no specific pathology (L). black bar=3 mm

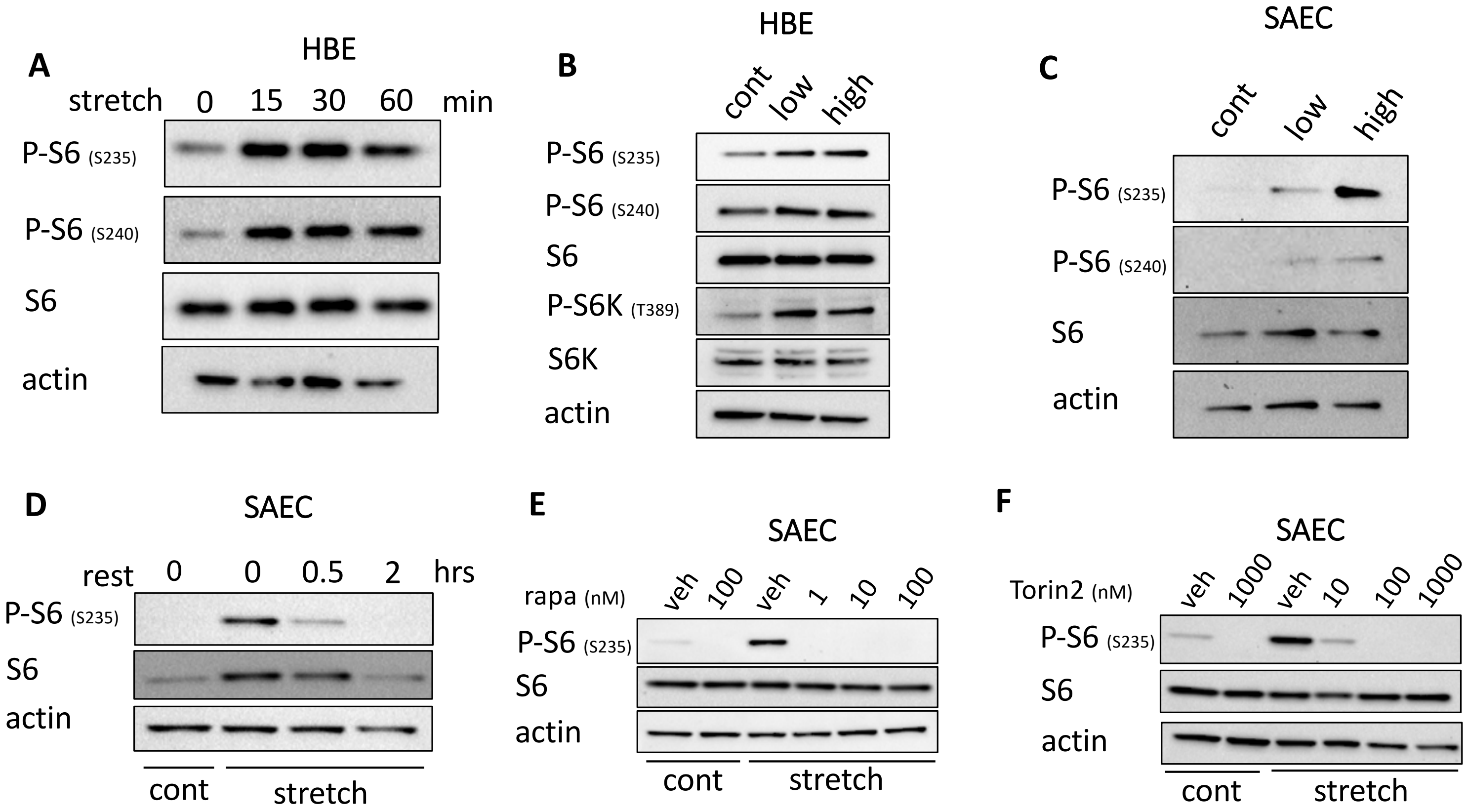


**Supplemental Figure 7. Volutrauma rapidly activates mTORC1 in a dose dependent fashion.** A) Human bronchial epithelial cells (HBE) were subjected to equibiaxial stretch (20%, 0.2 Hz) for varying amounts of time prior to immunoblotting for markers of mTORC1 activation (protein pooled from n=3 wells/lane). B) Human bronchial epithelial cells (HBEs) were subjected to 12% biaxial stretch (low) or 24% biaxial stretch (0.3 Hz) or static culture as control (cont) for 30 min prior to immunoblotting for markers of mTORC1 activation (protein pooled from n=6 wells/lane). C) Small airway epithelial cells (SAECs) were subjected to 12% biaxial stretch (low) or 24% biaxial stretch (0.3 Hz) for 4 hrs prior to immunoblotting for markers of mTORC1 activation (protein pooled from n=6 wells/lane). D) SAECs were subjected to injurious biaxial stretch (24% stretch, 0.5 Hz, 4 hrs) and then placed back into static culture (rest) for varying amounts of time prior to immunoblotting for markers of mTORC1 activation. (protein pooled from n=6 wells/lane) E) SAECs were subjected to in vitro volutrauma or static culture in the presence of increasing concentrations of rapamycin (rapa) for 4 hours prior to immunblotting for markers of mTORC1 activation (protein pooled from n=3 wells/lane) F) SAECs were subjected to in vitro volutrauma or static culture in the presence of increasing concentrations of Torin 2 for 4 hours prior to immunblotting for makers of mTORC1 activation (protein pooled from n=3 wells/lane)

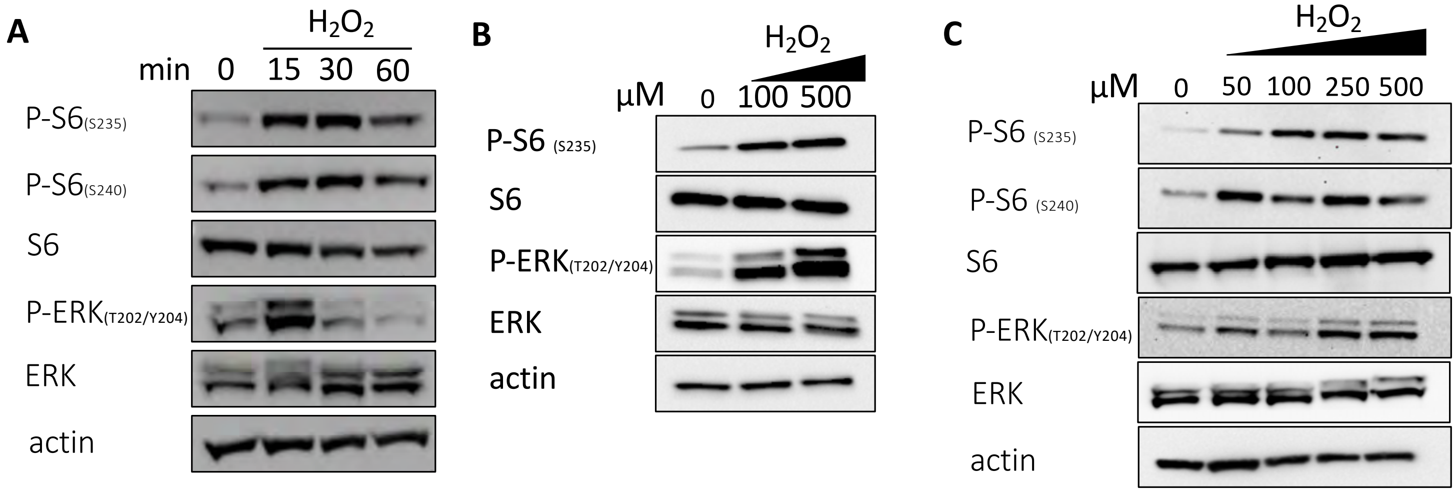


**Supplemental Figure 8. Hydrogen peroxide rapidly activates mTORC1 in airway epithelial cells in a dose dependent fashion.** A) Human bronchial epithelial cells (HBEs) were treated with 100 μM hydrogen peroxide ($H_2O_2$) for increasing amounts of time prior to immunoblotting for markers or ERK and mTORC1 activation. (protein from n=3 wells/lane) B) HBEs were treated with increasing doses of $H_2O_2$ for 30 min prior to immunoblotting for markers of ERK and mTORC1 activation. (protein from n=3 wells/lane) C) Small airway epithelial cells (SAECs) were treated with increasing doses of $H_2O_2$ for 30 min prior to immunoblotting for markers of ERK and mTORC1 activation. (protein from n=1 well/lane)

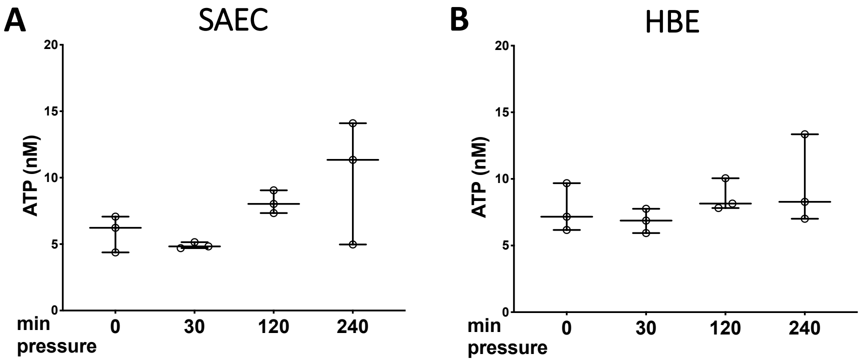


**Supplemental Figure 9. In vitro barotrauma does not increase extracellular ATP release in airway epithelial cells.** Small airway epithelial cells (SAEC, panel A) and human bronchial epithelial cells (HBE, panel B) were subjected to in vitro barotrauma (20 cm $H_2O$, 0.2 Hz) for varying amounts of time prior to measuring extracellular ATP. (n=3 wells/timepoint)
